# *Drosophila Prickle* Mutants Display Comorbid Neurological Phenotypes and Provide A Genetic Link Between Epilepsy and Autism Spectrum Disorder

**DOI:** 10.64898/2026.07.20.739611

**Authors:** Krishna M. Nukala, Brady Williquett, Anthony J. Lilienthal, Dakota M. Thompson, Josephine N. Massingham, Shu Hui Lye, Avery Yu, Bridget C. Lear, G. Gregory Neely, Stanislava Chtarbanova, J. Robert Manak

## Abstract

Epilepsy affects approximately 30% of individuals with autism spectrum disorder (ASD). Consistent with these observations, while *PRICKLE* mutations are primarily linked with epilepsy, there is an enrichment of pathogenic DNA sequence variants in *PRICKLE* genes carried by individuals with ASD. Nonetheless, a connection between *PRICKLE* function and ASD warrants further investigation. Here, we show that a seizure-prone *Drosophila prickle* mutant (*prickle-spiny-legs*, or *pk^sple^*) exhibits learning and memory deficits, increased pain sensitivity, both communication and social interaction difficulties, and restrictive repetitive grooming behaviors, all of which are strongly correlated with ASD, while a non-seizure prone *prickle* mutant (*prickle-prickle*, or *pk^pk^*) does not, thereby providing a direct genetic connection between epilepsy and ASD through *prickle*. Comparing headed versus headless *pk^sple^* mutants, we also show that the excessive grooming requires higher level cognitive processing from the brain. Finally, both *pk^sple^* and *pk^pk^* mutants exhibit circadian rhythm defects, another feature correlated with ASD, as well as distinct yet overlapping neurological anomalies in processes that include innate immune response, oxidative stress response, neuronal cell death, neurodegeneration, motor dysfunction and reduced lifespan, likely reflecting the unique isoform expression patterns observed in the developing CNS. Collectively, this study highlights the broadscale effects of *PRICKLE* mutations that extend beyond the primary clinical features of epilepsy to include several of the core features of ASD.

## 1. Introduction

In recent years, studies involving *Drosophila* models of neurological disease have gained traction, significantly contributing to our understanding of the underlying disease mechanisms and associated phenotypes (Bolus et al., 2020; Bonini & Fortini, 2003). Most of these studies have focused on 1) investigating human neurological disease genes conserved in flies for which mutants have been generated, 2) creating flies that express the pathogenic form of a human disease gene, or 3) engineering a human disease-causing variant into the conserved fly gene (Feany & Bender, 2000; Hotta & Benzer, 1970; Kroll et al., 2015; Lin et al., 1998; Lush et al., 1998; Min & Benzer, 1999). Genetically tractable flies offer an exceptional opportunity to study a broad range of phenotypes and pathologies associated with a disease gene *in vivo* compared to *in vitro* human cell-based assays which are unable to fully replicate the disease pathology (Marsh & Thompson, 2006). However, many studies of established animal models (including flies) of neurological disease have focused on the most evident disease pathology (i.e., seizures in animal epilepsy models) with less consideration of other potentially relevant disease comorbidities. For example, studies involving *Drosophila* sodium channel mutants with increased seizure susceptibility have focused on identification of gene mutations that specifically cause seizure-suppression, with the goal of elucidating relevant seizure circuitry pathways (Kroll et al., 2015; L. Parker et al., 2011; Louise Parker et al., 2011). While these studies have greatly contributed to our understanding of disease pathways, most neurological disorders in humans are complex thereby demonstrating a need to leverage disease models that can explore the compendium of neurological defects contributing to the overall disease pathogenesis.

The *prickle* (*pk*) gene, first identified in flies, encodes three transcript isoforms, *prickle-prickle* (*pk^pk^*), *prickle-spiny-legs* (*pk^sple^*), and *prickle-M* (*pk^M^*). *pk^pk^*and *pk^sple^* isoform-specific mutants have been the focus of the vast majority of studies, particularly given the critical role of these isoforms in establishing epithelial cell polarity during fly development (Gubb et al., 1999). In contrast, *pk^M^* has been understudied largely due to the lack of (up until recently) *pk^M^*-specific mutants (See Supplementary Figure 1; (Cho et al., 2022)). However, the role of *pk* in nervous system development and function has remained underexplored in spite of the observation that *prickle* shows some of its highest expression in the developing larval central nervous system (Brown et al., 2014). Work in our laboratory and others have shown that mutations in *PRICKLE* orthologs lead to an increase in seizure susceptibility in flies, fish and mice suggesting that the underlying mechanism governing epilepsy is conserved (Ehaideb et al., 2014; Ehaideb et al., 2016; Mei et al., 2013; Tao et al., 2011). Homozygous loss-of-function (LOF) mutations in the *pk^sple^* but not *pk^pk^* isoform lead to unprovoked, spontaneous myoclonus-like seizure activity and ataxia similar to those seen in human *PRICKLE* patients (Ehaideb et al., 2016), while homozygous LOF mutations in the *pk^pk^* isoform result in reduced viability of embryos (Ehaideb et al., 2014). Furthermore, mutations in *PRICKLE* orthologs have been found to affect migration of facial branchiomotor neurons in fish and mice (Mapp et al., 2011; Yang et al., 2014) as well as axonal advance of mushroom body (*pk^sple^*mutants) (Ng, 2012) and abdominal sensory neurons (*pk^pk^* mutants) (Mrkusich et al., 2011) in flies. Studies in mice have shown that *Prickle* genes promote formation and maintenance of glutamatergic synapses (Ban et al., 2021) as well as regulate axon number and axon initial segment (AIS) maturation by modulating microtubule bundling (Dorrego-Rivas et al., 2022). Further work in our laboratory connecting *prickle* and microtubule dynamics has shown that *pk* mutants affect microtubule-based vesicular transport in motor neuron axons whereby *pk^sple^* mutants show enhanced anterograde vesicle transport while homozygous *pk^pk^* mutants show reduced overall transport of vesicles in both directions, and that overexpression of a specific *pk* isoform (*pk^sple^*) can reverse axonal microtubule polarity in the motor neurons (Ehaideb et al., 2014). Additional work identified a significant increase in neuronal cell death and seizures with age in *pk^sple^* mutants that is driven by oxidative stress and the glial innate immune response (Nukala et al., 2023). Finally, a recent study has shown that *pk^M^*, previously thought to be exclusively expressed during embryonic stages, is also expressed in the developing *Drosophila* ommatidia although loss-of-function mutants were not found to exhibit PCP or overt behavioral phenotypes (Cho et al., 2022).

Autism spectrum disorder (ASD), a neurodevelopmental disorder, is characterized by a variety of clinical features including difficulties in communication and social interactions (e.g., trouble reading non-verbal cues, expressing needs), restrictive or repetitive behaviors (e.g., interests in a limited number of subjects, lining up objects, hand-flapping, body rocking, repetitive movements of the body), and altered responses to stimuli (e.g., the inability to differentiate the strengths of different stimuli) (Caldwell-Harris, 2021; Fuentes et al., 2021). In addition to these characteristics, ASD individuals can also present with a wide range of comorbidities including attention-deficit/hyperactivity disorder (ADHD), epilepsy, intellectual disability, neuroinflammation, neuropathies, sensory problems, and sleep disorders amongst others (Black et al., 2014; Kaye et al., 2024). Notably, nearly 30% of ASD patients are diagnosed with epilepsy (Viscidi et al., 2013) and our previous studies have shown that human *PRICKLE1* and *PRICKLE2* pathogenic mutations are enriched in individuals with ASD while mice carrying *Prickle1* and *Prickle2* gene disruptions exhibit ASD-like behaviors (Paemka et al., 2013; Sowers et al., 2013). Multiple studies have also revealed that individuals diagnosed with ASD can be hypersensitive (Failla et al., 2020; Hoffman et al., 2023; Qian et al., 2024) or insensitive (Chien et al., 2017; Dubois et al., 2020; Frundt et al., 2017) to different types of nociceptive stimuli, while disruptions in circadian rhythm patterns have been frequently reported in patients with ASD as well as epilepsy (Cohen et al., 2014; Jin et al., 2020; Pinato et al., 2019; Xavier, 2021), leading to disruptions in sleep patterns. The high co-occurrence of intellectual disability in individuals with ASD (with estimates ranging from 40-70%) impacts not only adaptive functioning such as social and communication skills but also learning, problem solving, and judgement (Buescher et al., 2014; Matson & Shoemaker, 2009; McKenzie et al., 2023). Together, these correlations encouraged us to assess whether *Drosophila pk* mutants display additional neurological comorbidities that could provide insights into the role *prickle* may have in bridging epilepsy and ASD.

In this study, we first show that *pk* transcript isoforms are expressed in the larval brain in distinct patterns, with *pk^sple^*and *pk^M^* isoforms exhibiting widespread, largely overlapping expression, while the *pk^pk^* isoform shows more localized expression. Second, we show that there are significant overlaps between the *pk^sple^* and *pk^pk^* mutant phenotypes, including upregulation of innate immune response and oxidative stress mitigator genes, increases in neuronal cell death, and reductions in lifespan. In contrast, we show several key differences in the phenotypes of these mutants (potentially due to the unique expression patterns of these isoforms), with an increased neurodegeneration index and double-strand DNA breaks only found in *pk^pk^* mutant brains, and increased seizure penetrance, ataxia, and locomotor defects specific to *pk^sple^* mutants. Third, we reveal that the seizure-prone homozygous *pk^sple^* but not *pk^pk^* mutants display significant learning and memory defects (features strongly correlated with ASD) which led us to explore whether *pk^sple^* mutants exhibited additional characteristics of ASD. Intriguingly and in addition to the learning deficits, *pk^sple^* mutants (but not *pk^pk^*mutants) had a greater sensitivity to thermal pain, displayed social interaction difficulties, and exhibited restrictive and repetitive behavior in the form of enhanced grooming activity, while both mutants showed circadian rhythm defects reflecting sleep disturbances, all comorbidities associated with ASD. Together these data provide compelling evidence connecting epilepsy and ASD through *prickle*.

## 2. Methods

### 2.1. Fly Husbandry and Fly Lines

All flies (*Drosophila melanogaster*) were reared in a 25°C incubator on a 12h light/dark cycle using standard cornmeal-molasses medium food and for histology and BBB assays, Nutri-Fly^®^ Bloomington formulation food (Genesee Scientific, Cat #: 66-113) was used. Stocks used were *Oregon-R* (obtained from Chung Fang Wu’s laboratory, University of Iowa), *w^1118^* (obtained from Andy Frank’s laboratory, University of Iowa), *Canton-S* (obtained from Josh Dubnau’s laboratory, Stony Brook University), *pk^pk^* and *pk^sple^* which were outcrossed into *Oregon-R*, *w^1118^* and *Canton-S* backgrounds for a minimum of 7 generations to minimize genetic background issues, *UAS-RelishRNAi* (BDSC: 33661), *repo-Gal4* (BDSC: 7415) and *dTrpA1^ins^* (obtained from Paul Garrity, Brandeis University).

### 2.2. HCR Fluorescence in situ Hybridization Assay

Probe sets and amplification hairpins for the detection of the different *prickle* transcript isoforms in the *Drosophila* 3^rd^ instar larval brain were designed by Molecular Instruments^TM^ (pk-M-B2/B2-Alexa Fluor® 647, pk-pk-B3/B3-Alexa Fluor® 546, and pk-sple-B4/B4-Alexa Fluor® 488). The sequences for the probes are listed in Supplementary Figure 2. Hybridization buffer, amplification buffer, and wash buffer were obtained from Molecular Instruments^TM^. Third instar larval brains were dissected in 1x nuclease-free PBS and then fixed and prepared as described (Young et al., 2020; Younger et al., 2022). The brains were mounted using Vectashield H1000 and analyzed with a Leica TCS SP8 confocal microscope using a 10x dry objective. Images were acquired using Leica’s LAS X Software.

### 2.3. Immunohistochemistry and Confocal Microscopy

Adult fly brains were dissected in 1X PBS and fixed in 4% paraformaldehyde in 1X PBS for 40 minutes, washed for 5×10 minutes in 0.03% PBS-Triton. All brains were incubated with blocking solution (0.1% PBS-Triton-BSA) for 40 minutes followed by overnight incubation at 4°C with primary antibodies. Samples were washed for 6×10 minutes with 0.03% PBS-Triton followed by incubation for 40 minutes at room temperature in blocking solution. Next, samples were incubated for 2 hours at room temperature with secondary antibodies diluted in blocking solution. After 6 successive 10 minute washes with 0.03% PBS-Triton, brains were mounted in Vectashield^®^ with DAPI mounting medium (Vector Laboratories). Brains were analysed under Leica TCS SP5 confocal microscope using a 20X dry objective or a 63X water objective and images were acquired using Lecia’s LAS AF software. The following primary antibodies were used: Rabbit anti-Dcp1 (1:100) (Cell Signaling Technologies), Rat-Elav-7E8A10 anti-elav (1:200) was deposited to the DSHB by Rubin, G.M. (DSHB Hybridoma Product Rat-Elav-7E8A10 anti-elav), Mouse-UNC93-5.2.1 anti-H2AV (1:500) was deposited to the DSHB by Hawley, R.S. (DSHB Hybridoma Product UNC93-5.2.1). The following secondary antibodies were used: Alexa Fluor^®^ 488-conjugated goat anti-rabbit (1:200 for anti-Dcp1) (Invitrogen), Alexa Fluor^®^ 488-conjugated goat anti-mouse (1:500 for anti-H2AV) (Invitrogen), Alexa Fluor^®^ 568-conjugated goat anti rat (1:300 for anti-elav) (Invitrogen).

### 2.4. Image Processing and Quantification

ImageJ (Schneider et al., 2012) was used for quantification and image processing. 30-40um maximum projection images of the adult mid-brain were used to quantify the number of Dcp-1 and H2AV positive puncta. Background noise was removed using background subtraction (rolling = 5) and Gaussian blur filter (sigma= 0.5-1) or despeckling. Next, the images were manually thresholded and the apoptotic or H2AV positive puncta (defined as fluorescent particles greater than 0.2 micron^2^ after background noise removal and thresholding) were counted.

For Dcp-1 and H2AV quantification, the number of Dcp-1 and H2AV positive puncta were assumed to be 100% in the control for normalization purposes.

For HCR quantification, we used LAS X to take 20x magnification stacks with 1 um slices in each of the following brain regions: both optic discs (ODs) and laminas, the anterior and posterior regions of both medullas, the central brain (CB) and the anterior subesophageal zone (SEZ), the posterior SEZ and the thoracic ventral nerve cord (VNC), and the abdominal VNC. Image processing was conducted in ImageJ. The threshold was determined manually and despeckled, then watershed analysis was applied. The stack was then converted to individual images and the number of puncta expressing each isoform in each image was counted using the “Analyze Particles” tool. To identify puncta expressing both *pk^pk^* and *pk^sple^*isoforms, we then used the “Image Calculator” tool to identify the pixels with both *pk^pk^* and *pk^sple^* expression and the resulting puncta were counted using the “Analyze Particles” tool.

### 2.5. Histology and Neurodegeneration

Flies of indicated genotypes were collected at 0-3 or 0-4 days after eclosion and aged at 25°C for up to 5- (range 3-8 days), 15- (range 14-18 days) or 30- (range 26-32 days) day old. At the desired age, fly heads were severed using a razorblade and placed in Carnoy’s fixative (100% Ethanol:chloroform:acetic acid at 6:3:1) over night at 4°C. The fixation solution was replaced with 70% Ethanol the next day and the head paraffin embedding procedure, head sectioning (at 5 µm) and hematoxylin and eosin (H&E) staining were followed as described in (Gevedon et al., 2019). The presence of neurodegeneration was indicated by the appearance of holes in the brain neuropil and was scored blindly for each genotype using the 6-level (scale of 0,1,2,3,4 and 5) neurodegeneration index score as described in (Cao et al., 2013). Images were acquired using a Camera-equipped Nikon Eclipse E100 light microscope using the Nikon NIS-Elements image acquisition software. Acquired images were next processed using Adobe Creative Cloud Photoshop software for proper orientation in respective figures.

### 2.6. Blood Brain Barrier Assay

Blood eye barrier (BEB) integrity was used as a proxy to determine blood brain barrier (BBB) integrity (DeSalvo et al., 2011; Pinsonneault et al., 2011). Female flies (3-10 days old) of each genotype were individually injected under CO_2_-anesthesia with 69nL of 50mg/mL of 10,000 MW Alexa Fluor^®^ 488-Dextran (Invitrogen^TM^ catalogue # D22910) using a glass capillary (Drummond 3-000-203-G/X) mounted on a Nanoject II Auto nanoliter injector (Drummond Scientific). Injected flies were incubated for 2 hours at 25°C, following which they were anesthetized on ice, and placed under a fluorescent stereoscope to score BBB integrity. The presence of a fluorescent hemolymph exclusion line or diffused eye fluorescence was used to score intact and disrupted BBB, respectively. Representative images of eyes were acquired using a DSLR camera mounted on the stereoscope.

### 2.7. Video-Tracking Locomotor and Climbing Assay

The video-tracking locomotor assay using pySolo was performed as described in (Ehaideb et al., 2016) with a few modifications. All the flies assayed were aged for 5, 15 and 30 days post-eclosion (dpe) and were separated by sex a day prior to when the assay was performed and allowed to acclimate in an environmentally controlled chamber (Chamber Temperature: 25°C, Relative Humidity: 60%) in which the assay was performed. All videos taken were subjected to a single blind analysis to avoid bias. Total activity (in pixels) and median turning angle was determined using pySolo.

A negative geotaxis (climbing) assay was used to quantify locomotion defects in flies aged for 5, 15, and 30 dpe. At least 12 hours before the start of the assay, groups of 9-10 adult flies of each genotype were anesthetized using carbon dioxide and separated by sex. To limit the effects of circadian rhythm alterations on the locomotor ability, all assays were conducted at the same time of day and all genotypes were tested in parallel. The flies were tapped to the bottom of an empty glass vial and videotaped for 10 seconds for three independent trials. For each trial, the climbing pass rate was quantified as the percentage of flies climbing 2 cm from the bottom of the vial and the overall climbing pass rate across the three trials was calculated by averaging the three independent trials.

### 2.8. Lifespan Analysis

Flies were collected at 1 dpe and grown at 25°C in separate vials. Flies were transferred to fresh vials every other day and number of flies alive in each vial was quantified every day until the death of the fly.

### 2.9. Gene Expression and Gene Ontology Analysis

Flies were aged to 7 days before dissecting their brains for microarray analysis which was performed as described (Nukala et al., 2023; Santana et al., 2020). Two biological replicates with 4 technical replicates were used to perform the gene expression analysis. Genes were considered differentially expressed if showing an FDR-adjusted *p* value of less than 0.05 and a fold change value of ≥ 1.5 (upregulated) or ≤ 0.66 (downregulated). Gene ontology (GO) analysis was performed using DAVID (Huang et al., 2009a, 2009b) by inputting the list of differentially expressed genes and a list of significantly enriched GO terms were obtained.

### 2.10. Thermal Nociception Assay

Flies were raised on a standard diet (cornmeal, yeast, molasses, and agar) at 25°C, 65% humidity and 12-h:12-h light:dark cycle for 7 days before undergoing the heat nociception assay as described in (Khuong et al., 2019). Animals were placed onto a hot plate and covered with an arena made from a modified 35mm petri dish. Temperature was increased from 25 to 50°C over 195 seconds. Tracking for velocity analysis was performed using Tracktor software (Sridhar et al., 2019).

### 2.11. Olfactory Aversion Assay

For this assay, 3-octanol (OCT) and 4-methylcyclohexanol (MCH) scents (attractive at low concentrations and aversive at high concentrations (Wang et al., 2003)) were used to train 50 3–5-day old adult flies after placing them into the T-maze apparatus (without electric shock). After providing 1 minute of air flow to acclimate to the testing chamber, these flies were then exposed to varying (0%, 10% 20% 40%) 3-octanol on one side and 1:500 4-methylcyclohexanol on the other. Performance index was calculated based on the number of flies which moved away from the branch containing 3-octanol. Both odors were administered at ∼80 mL min-1.

### 2.12. Shock Avoidance/Aversive Olfactory Conditioning Assays

Aversive Pavlovian olfactory association was assessed by training flies in a T-maze apparatus with a Pavlovian conditioning procedure (Li et al., 2013; Tully & Quinn, 1985). At least 36 hours prior to the start of the assay, 50 flies were separated into vials (50%/50% male/female). All assays were conducted on 7-10 dpe flies in dim red light at 25°C and ∼60% humidity. On the day of the assay, the white lights were turned off 2 hours prior to the start of the assay and the red light was turned on 1 hour before the assay to allow the flies to acclimate to the conditions of the assay. Both assays were conducted reciprocally to eliminate side and tube bias, and all genotypes were assayed in parallel. The shocks for both shock avoidance and aversive conditioning assays were delivered by a Grass S48 stimulator at 90 V at an interval of one 2-second shock per 5 seconds. To assay shock avoidance, flies were allowed to acclimate to the T-maze for 2 minutes and then transferred to the choice point with both arms lined with an electrical grid, one electrified and the other not. The flies were allowed to choose between the two chambers for 1 minute and then separated based on their choice and counted. For aversive conditioning assays, the shocks were associated with either MCH (Sigma, diluted 1/1000 in light paraffin oil vol/vol) or OCT (Sigma, diluted 0.5/1000 in light paraffin oil vol/vol) at ∼80 mL min-1. For a single training session, the flies were exposed to the scents sequentially with 1 minute of odor paired with an electric shock followed by one minute of the second odor without the shock with 45 seconds of rest in between. After exposure to the second scent, the flies were given an additional 45 seconds of rest before they were transferred to the choice point. At the choice point, the flies were given 2 minutes to choose between tubes containing one of the two scents and then separated based off of their choice. To minimize the potential for scent bias, the scent associated with the shock was alternated in consecutive assays. The results were separated by sex and quantified using the performance index (P.I.) metric as described (Tomchik & Davis, 2009). The calculation for quantifying olfactory conditioning is shown below, but the calculation for assaying shock avoidance was done using the same method. In short, for each reciprocal assay, the number of flies making the incorrect choice (shock-associated) were subtracted from the number of flies making the correct choice (no shock) and then divided by the total number of flies counted. The overall P.I. was then calculated by finding the average PI of the two reciprocal assays. Flies that failed to make a choice were not considered in this calculation.

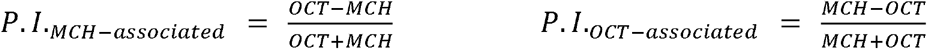

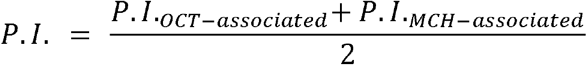

### 2.13. Circadian Rhythm Assay

All assays were performed at 25°C. Locomotor activity levels of adult male flies were measured for 5 days in a 12h light:12h dark (LD) conditions, followed by 7 days of exposure to constant darkness (DD), using the *Drosophila* Activity Monitor system (Trikinetics). For DD rhythmicity analysis, chi-squared periodogram analyses were performed on individual flies over 7 days using ClockLab analysis software, as previously described in (Moose et al., 2017). Flies were considered rhythmic if chi-squared power was greater than or equal to 10 above significance (Lear et al., 2005). To produce LD activity profiles, activity levels of individual flies were normalized and averaged within genotypes over the last 4 days of LD conditions. LD and daily activity profiles were generated using the Excel-based program Counting Macro (Pfeiffenberger et al., 2010). Morning and evening anticipation values were calculated in individual flies over the last 3 days of LD using the following formula: (total activity 3 hours before light transition)/ (total activity 6hr before light transition) – 0.5, as previously described (Seluzicki et al., 2014).

### 2.14. Electroretinogram

To perform ERG recordings, adult flies were anesthetized, and their heads were fixed in place on a copper mesh using wax. A grounding (reference) electrode was inserted into the proboscis and the recording electrode was inserted into the cornea of the eye. Flies were allowed to acclimate to the dark for at least 5 minutes prior to being exposed to a train of 10, 1-second flashes of white light pulses. Retinal responses were recorded and analyzed using Labchart software.

### 2.15. Social Isolation Assay

To assess levels of social interaction within a defined space, a transparent acrylic circular socialization chamber (SC; outer edge Ø = 95.25 mm; inner edge Ø = 88.9 mm; chamber height = 1.905 mm) was created (University of Iowa Biology Engineering Shop). 5 males and 5 females (all aged to 7-10 dpe) were anaesthetized by cold shock for 75 seconds and then transferred to a large petri dish where they were trapped within the SC for recording. After transfer to the SC, flies were allowed to recover from the cold shock for 8 minutes and then one minute video recordings of the flies captured at 30 frames per second were taken at 8, 11, and 14 minutes. The positions of the flies were recorded using ivTools (https://opensource.cit-ec.de/projects/ivtools) and every tenth frame over the course of the full video was used to calculate the distance of each fly relative to each other fly (interindividual distance) using Matlab_R2024b (Mathworks inc.). 30 total flies for each genotype were assayed, and all genotypes were tested in series on the same day.

### 2.16. Male Courtship Behavioral Assay

In order to assess social communication through measurement of male courtship behavior, naive males from each genotype (controls, *pk^sple^*, *pk^pk^*; 20 flies each) and virgin control females (20 flies/each genotype analyzed) were first aged in isolation in vials containing standard *Drosophila* medium for a total of 5 dpe. One male and one control female were then placed together in a circular mating chamber by aspiration within the ZT0-ZT3 time window (Dey et al., 2024). After a five-minute adjustment period, the flies were recorded for 15 minutes at 30 frames per second. Courtship behavior was analyzed in a blinded fashion using Video Viewer in MATLAB to manually score the number of frames spent pursuing the female, adjusting their orientation relative to the female, wing flickering, licking, attempting to copulate, or copulating. The number of frames spent courting (a summation of all courtship parameters) out of the 27,000 total frames generated the courtship index. Courtship latency was determined by calculating the time elapsed before the male engaged in any courtship behaviors, and copulation attempts represents the total number of attempts to copulate over the 27,000 frames. Males copulating for the entire video duration were excluded from copulation attempt calculations but were included in courtship index and latency calculations.

### 2.17. Grooming Behavior Assay

To quantify repetitive behavior, spontaneous grooming was analyzed using 5 dpe flies placed in circular chambers (acclimated to an environmentally controlled room; 25 °C, relative humidity 60%) using high-resolution videography (five minutes, 9,000 frames total) as previously described (Nukala et al., 2023). Videos were manually scored for grooming behavior by an observer blind to the genotype using the Video Viewer tool in MATLAB R2024a. A grooming bout was defined as any instance of continuous grooming lasting greater than or equal to 4 frames (∼130 milliseconds) of continuous grooming. Start and stop frames were recorded for each grooming bout and then classified by the type of grooming observed as described: front legs, head/eyes, back legs, abdomen, and wings (Szebenyi, 1969). Thorax grooming was not recorded due to its rare occurrence in all genotypes tested. The grooming bouts were then further classified into anterior or posterior cleaning modules based on which pair of legs (e.g., front or hind) was involved in grooming (Mueller et al., 2019). The percent time spent grooming, grooming bout number (B#), and median grooming bout duration (BD) were then calculated for each fly. Statistical significance was determined using a one-way ANOVA with Tukey’s multiple comparisons test.

### 2.18. Decapitated Fly Grooming Analysis

To determine whether the brain has a role in excessive grooming behavior in *pk^sple^* mutants, grooming assays were conducted on decapitated 15 dpe *pk^sple^* mutants with age-matched, head-intact control and *pk^sple^*mutants. Flies were separated by sex using CO_2_ anesthesia at least 12 hours prior to the start of the assay and allowed to acclimate to a temperature/humidity-controlled room (25°C, 60% relative humidity). To limit the effects of anesthesia on grooming, flies were not anesthetized prior to decapitation. Instead, flies were held in place by vacuum within a finely cut 2-mm pore of a P1000 pipette tip covered with Flystuff® *Drosophila* mesh (Genessee Scientific). Flies were then either decapitated with a 2.5 mm Vannas Spring scissors (Fine Science Tools) or immediately transferred to an empty vial (head-intact controls) and allowed to acclimate to the temperature and humidity-controlled room for 15 minutes before being assayed. Before recording, flies were transferred by mouth aspirator to an analysis chamber and allowed to acclimate to the lightbox for 5 minutes, after which the decapitated flies were positioned upright in the well immediately before recording started. Only decapitated *pk^sple^* mutant flies that showed a vigorous righting response when pushed over following the grooming assay were analyzed for spontaneous grooming for a duration of 5 minutes (Yellman et al., 1997).

### 2.19. Statistical Analysis

Statistical analysis was performed using GraphPad Prism 10. All data were assessed for Gaussian distribution using the Shapiro-Wilk’s normality test and analyzed with a two-tailed unpaired Student’s t test, Mann-Whitney U test, One-way ANOVA, Two-way ANOVA and Kruskal-Wallis test. Error bars for all data represent the mean ± SEM. Two-tailed *p* values of < 0.05 were considered the cutoff for statistical significance.

## 3. Results

### 3.1. *pk^sple^* and *pk^M^* transcripts show widespread, overlapping brain expression while *pk^pk^* expression is widespread but more focalized

The distinct neurological phenotypes observed in mutants of different *prickle* transcript isoforms (Ehaideb et al., 2014; Ehaideb et al., 2016; Nukala et al., 2023) implies that the different isoforms may be expressed in distinct patterns. In order to assess expression of all three *prickle* transcript isoforms (*pk^pk^, pk^sple^,pk^M^*), we performed an *in situ* hybridization analysis using the Hybridization Chain Reaction (HCR) fluorescence in situ hybridization assay (Young et al., 2020; Younger et al., 2022). In the larval thoracic ventral nerve cord (VNC), the subesophageal zone (SEZ), and the central brain (CB) region (Figure 1A), *pk^sple^*is primarily expressed in areas of the brain containing a high percentage of type I neuroblasts (NBs; Figure 1B), neural stem cells that asymmetrically divide into NBs and ganglion mother cells (GMCs; the immediate precursor to neurons and glia in the developing brain) (Homem & Knoblich, 2012), and immature (or recently matured) neurons, in a pattern that is similar to that of *prospero* which itself is expressed at lower levels in neuroblasts but is upregulated in neurons (Moraru et al., 2012; Samuels et al., 2020). In the developing optic lobe (OL), *pk^sple^*exhibits high levels of expression on the medial side of the developing medulla (Me) and the portion of the lamina (La) on the lateral side of the laminar furrow (LF). Both regions are populated by NBs, with type I NBs making up the medial medulla and lamina precursor cells (LPCs) in the lateral lamina, a class of type I NBs unique to the lamina (Egger et al., 2007). There is lower-level expression of *pk^sple^* in the lateral medulla and medial lamina, which make up the two arms of the outer proliferation center (OPC), a layer of neuroepithelial (NE) cells projecting inward from the surface of the developing OL (Supplementary Figure 3A). In the thoracic VNC, expression of the *pk^sple^* isoform mirrors that of the central brain, with high levels of *pk^sple^* found in the type I NBs and neurons of the VNC. Overall, these findings indicate that the *pk^sple^* isoform is highly expressed primarily in type I NBs and immature or recently matured neurons throughout the larval brain and VNC, but not in the NE of the developing optic lobe.

**Figure 1.**
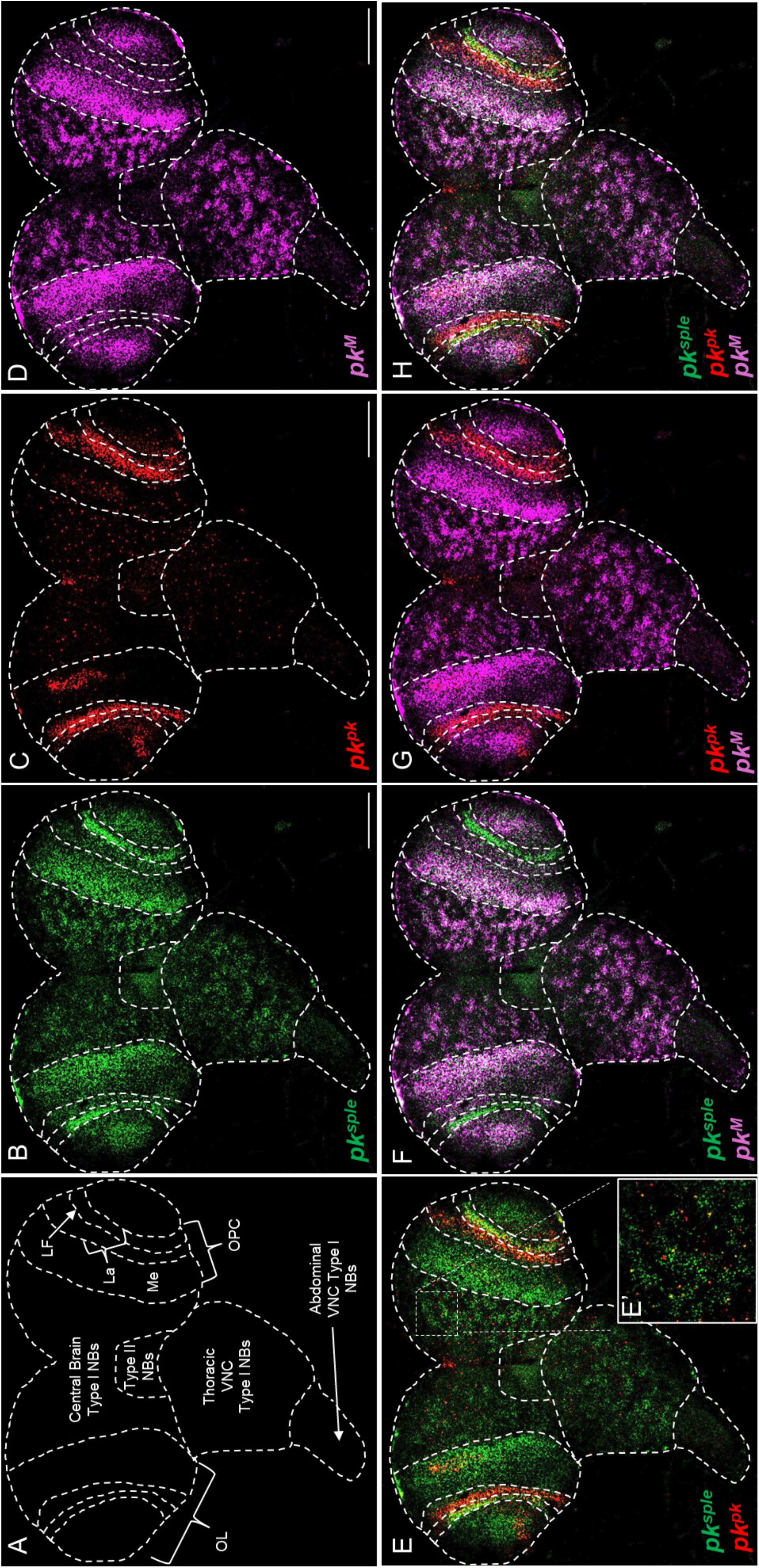
***pk^M^* and *pk^sple^* isoforms are highly expressed throughout the wild-type third instar larval brain, while the expression pattern of the *pk^pk^* isoform is more discrete.** A, Schematic depicting the anatomy of the Drosophila third-instar larval brain. **B-D,** Single channel images depicting the expression patterns of **B,** sple-B4, **C,** pk-B3, and **D,** pkM-B2 probes visualized with B4-AlexaFluor®488, B3-AlexaFluor®546 and B2-AlexaFluor®647 respectively, and amplifiers in multiplexed *in situ* HCR in a single slice through the brain. Scale bar =100 μm. **E-G,** Two channel overlaid images depicting E, *pk^sple^*-*pk^pk^*, F, *pk^sple^*-*pk^M^*, and G, *pk^pk^-pk^M^*expression patterns. **E’,** Inset image of the top right central brain region of the brain depicting colocalization of *pk^sple^* and *pk^pk^*in nearly all cells expressing the *pk^pk^* isoform. **H**, Three channel overlay representing expression of all three isoforms within the same brain. Abbreviations: La: Lamina; LF: Laminar Furrow; Me: Medulla; NBs: neuroblasts; OL: Optic Lobe; OPC: Outer Proliferation Center; VNC; Ventral Nerve Cord.

The *pk^pk^* isoform exhibits a more specific, localized expression profile than *pk^sple^* (Figure 1C and Supplementary Figure 3B). Throughout the larval brain, the number of cells expressing only *pk^pk^*is significantly lower than those expressing only *pk^sple^* (Supplementary Figures 3A and 3B). In the VNC and CB, *pk^pk^* is only observed in isolated cells, not in large cell clusters as observed for *pk^sple^*. (Figure 1E). Upon closer examination of these transcripts in the CB, the *pk^pk^-*expressing cells are frequently found within larger clusters of *pk^sple^* expressing cells (Figure 1E’, Supplementary Figure 3C and 3D). In the OL, the level of *pk^pk^* expression is much higher than in the CB. Like *pk^sple^*, *pk^pk^* is expressed in the lateral portion of the La, but unlike *pk^sple^*, the *pk^pk^* transcript is also highly expressed in the NE of the medial La and lateral Me (Supplementary Figure 3A-D). The presence of both isoforms within the lateral La highlights a potential fate switch mechanism in laminar development, in which co-expression of *pk^sple^* and *pk^pk^* signals that the NE of the medial La will differentiate into LPCs. Taken together, these findings indicate that while *pk^pk^*appears to be expressed primarily in immature NE cells, *pk^sple^*activation in *pk^pk^* expressing cells may mark the initiation of transition from NE cells to type I NBs and neurons.

Expression of the *pk^M^* transcript closely resembles that of *pk^sple^* (Figures 1D and 1F). In the CB, VNC, and medial Me of the OL, the expression pattern of *pk^M^* is identical to *pk^sple^*, with nearly all cells expressing *pk^M^* co-expressing *pk^sple^*thus indicating that the type I NBs and neurons of the CB express both isoforms. In contrast to *pk^sple^, pk^M^* is not expressed in the lateral La. Within the OL, expression of *pk^M^* and *pk^pk^* is anti-correlated: where *pk^M^* expression is observed, there is no *pk^pk^* expression, and vice versa (Figure 1G). Collectively, these data indicate that *pk^sple^* and *pk^M^* transcript isoforms have largely overlapping and broadscale patterns in the larval brain, while *pk^pk^* shows a more localized, albeit widespread, pattern of expression (Figure 1H) thereby suggesting that the different neurological phenotypes observed between the *pk^sple^* and *pk^pk^* mutants ((Ehaideb et al., 2014; Ehaideb et al., 2016; Nukala et al., 2023) and below) may be in part due to these expression differences. Given the absence of PCP or overt behavioral phenotypes associated with a recently generated *pk^M^* mutant (Cho et al., 2022), this study focuses on phenotypic differences between the *pk^sple^* and *pk^pk^* mutants.

### 3.2. Similar to *pk^sple^* mutants, innate immune response genes are significantly upregulated in *pk^pk^* mutants

In an effort to identify gene pathways potentially connecting the reduced viability, larval microtubule vesicle transport defects, and abdominal sensory neuron axon extension defects reported in *pk^pk^* mutants (Ehaideb et al., 2014; Mrkusich et al., 2011), we compared microarray data of 7-10 day old homozygous *pk^pk^*mutant brains to a previously published microarray dataset (GEO: GSE154686) of control brains processed under identical experimental conditions, and performed gene ontology (GO) analysis (Figure 2A; (Nukala et al., 2023)). By employing moderate fold-change cutoffs (>1.5, <0.66), we identified a total of 716 differentially regulated genes (adjusted p < 0.05; 323 upregulated, 393 downregulated) in homozygous *pk^pk^* mutant brains. Using the Database for Annotation, Visualization and Integrated Discovery (DAVID) functional annotation bioinformatics tools (Huang et al., 2009b), genes associated with the immune response pathway (GO enrichment score: 10.6) was identified as a top hit, similar to what we observed for homozygous *pk^sple^*mutants (Nukala et al., 2023). We also identified several categories of genes connected to cytolysis pathway, heat response, and olfactory behavior. Among the immune response pathway GO category were 25 significantly upregulated genes including those encoding numerous anti-microbial peptides (*Attacin-A*, *Attacin-C*, *Cecropin-A1*, *Cecropin-A2, Cecropin C, Drosocin, Metchnikowin, Diptericin*) and *IM18*. Interestingly, we found that *Attacin-D* and *IM-23* were significantly upregulated in *pk^pk^* but not in *pk^sple^* mutant brains (Figure 2B). Among the heat response GO category (enrichment score: 2.2), we found multiple upregulated *Turandot* stress response genes of the JAK-STAT pathway (Ekengren & Hultmark, 2001; Myllymaki & Ramet, 2014). In contrast to *pk^sple^*mutant brains, oxidation-reduction (redox) pathway genes in *pk^pk^*mutant brains were not highly enriched (enrichment score: 0.85), with only 8 versus 41 differentially regulated redox genes for *pk^pk^* mutant and *pk^sple^* mutant brains, respectively. Among the 8 differentially regulated redox genes in *pk^pk^* mutant brains, we identified several encoding cytochrome P450 enzymes such as *Cyp6a2, Cyp4e3, Cyp6a14* and *Cyp4p2* implicated in mitigating oxidative stress and detoxification processes (Hrycay & Bandiera, 2015) also upregulated in *pk^sple^* mutant brains. In addition, several key redox genes encoding mitochondrial NADH dehydrogenase, catalase, aldo-keto reductase, NADPH oxidase and glutathione transferases were not upregulated in *pk^pk^* mutant brains compared to *pk^sple^* mutant brains (Figure 2C).

**Figure 2.**
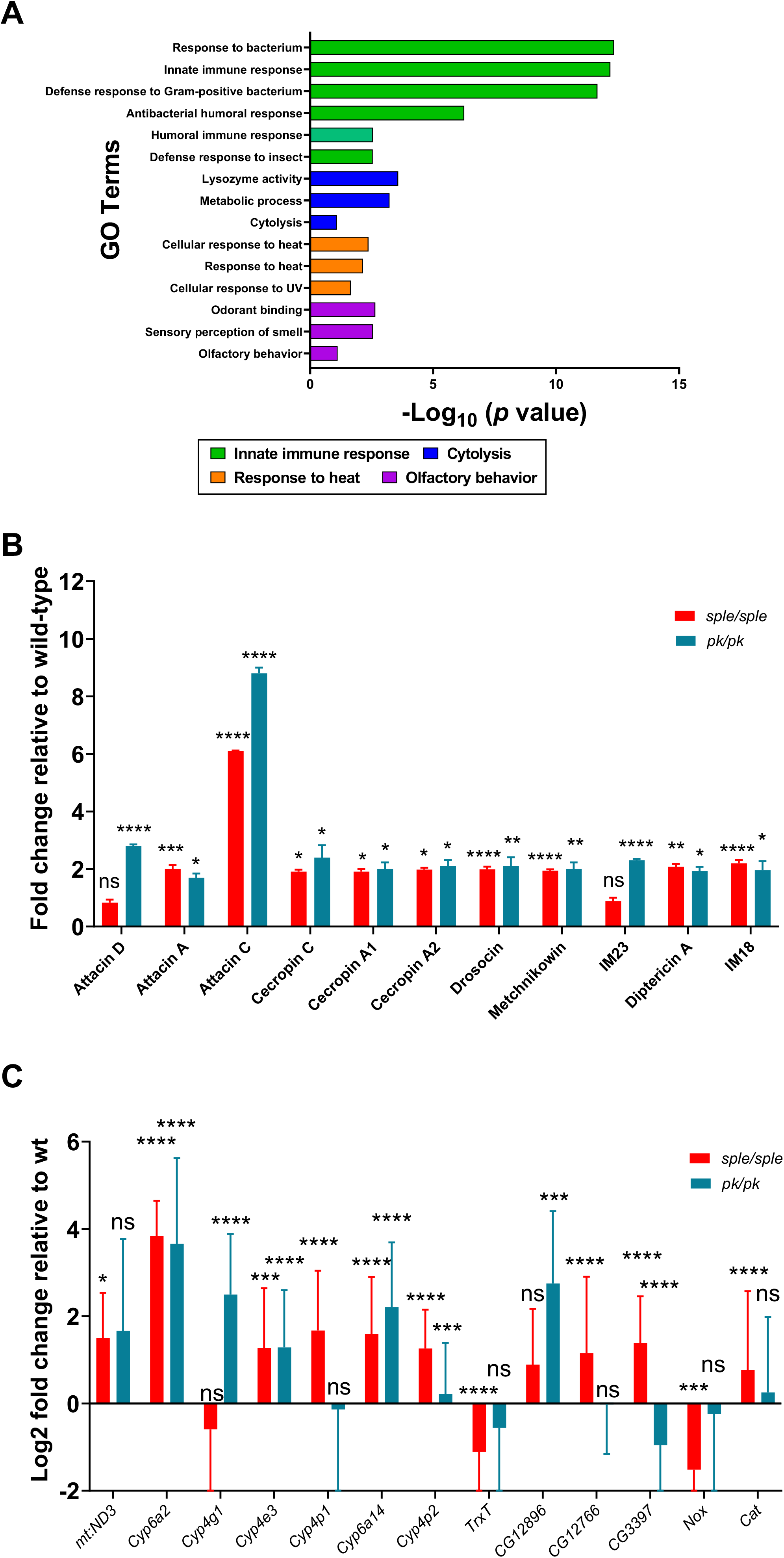
Homozygous *pk^pk^*mutants show significant increase in innate immune response gene expression. A, Gene ontology terms based on enrichment scores and are listed in descending order: Innate immune response, Cytolysis, Heat response and Olfactory behavior. **B,** Fold change values of specific AMP genes enriched in the microarray analysis of 7-10 days old *pk/pk* and *sple/sple* mutant brains relative to control. Majority of AMPs similar to what was observed in *sple/sple* mutants show a significant upregulation. Data are Mean ± SEM analyzed by Fisher’s exact test. **C,** Log_2_ fold change values of oxidative stress pathway genes differentially expressed in the brains of 7-10 days old *pk/pk* and *sple/sple* mutant brains relative to control. Several redox pathway genes were not significantly enriched or differentially expressed in *pk/pk* mutant brains when compared to *sple/sple* mutant brains except for genes encoding Cytochrome P450 enzymes, a majority of which are significantly upregulated in both *pk/pk* and *sple/sple* mutant brains. Data are Mean ± SEM analyzed by Fisher’s exact test. For all data, *p ≤ 0.05, **p ≤ 0.01 and ****p ≤ 0.0001. Note: Control and *sple/sple* microarray data were obtained from a previous publication (GEO: GSE154686) to facilitate comparison with *pk/pk* data (Nukala et al., 2023).

To determine whether the increase in the innate immune response in homozygous *pk^pk^* mutant brains is brain-derived, we performed a blood eye barrier (BEB, a proxy for blood brain barrier, BBB) integrity assay using injected fluorescently-labeled dextran in both homozygous *pk^pk^* and *pk^sple^*mutants compared to controls (Nukala et al., 2023). Notably, we failed to observe diffused fluorescence in the eyes of any injected *pk^pk^*or *pk^sple^* mutants, demonstrating that the BEB/BBB integrity is not compromised in either *pk* mutation (Supplementary Figure 4). Collectively these data demonstrate that, similar to *pk^sple^* and other fly neurological mutants (Cao et al., 2013; Petersen et al., 2013; Petersen et al., 2012), *pk^pk^* mutants (while not seizure prone) experience a significant upregulation of *AMPs* in the brain, yet show a reduced redox transcriptional response when compared to *pk^sple^* mutants. These data are consistent with the hypothesis that seizure activity drives increased oxidative stress, or vice versa (Fabisiak & Patel, 2022; Puttachary et al., 2015).

### 3.3. *pk^pk^* mutants show evidence of pronounced neuronal cell death, DNA damage and reduction in lifespan

Since upregulation of AMPs in the brain is known to lead to an increase in neuronal cell death and shortened lifespan, including in *pk^sple^*mutants (Cao et al., 2013; Kounatidis et al., 2017; Nukala et al., 2023; Petersen et al., 2012), we assessed whether this was also the case for *pk^pk^*mutants. Using an anti-Dcp-1 antibody (a marker for apoptotic cells), we observed a significant increase in the percentage of cleaved Dcp-1 positive puncta in homozygous *pk^pk^* mutant brains at all time points (5, 15 and 30 days post-eclosion, dpe) compared to age-matched controls. We also observed that *pk^pk^*mutant brains showed a significantly higher percentage of cleaved Dcp-1-positive puncta at 15 and 30 dpe compared to age-matched *pk^sple^*mutant brains (Figure 3A-D). This observation is supported by paraffin sectioning with H&E staining whereby *pk^pk^* mutants show a significantly higher neurodegeneration index at all timepoints examined compared to controls, and a significantly higher neurodegeneration index at 15 and 30 dpe when compared to *pk^sple^* mutant brains (Figure 3E-H). Taken together, these data demonstrate that *pk^pk^* mutant brains show evidence of sustained and widespread neuronal cell death and widespread vacuolization across adulthood that exceeds the neuronal cell death observed in *pk^sple^*mutant brains despite the absence of seizure activity in *pk^pk^*mutants.

**Figure 3.**
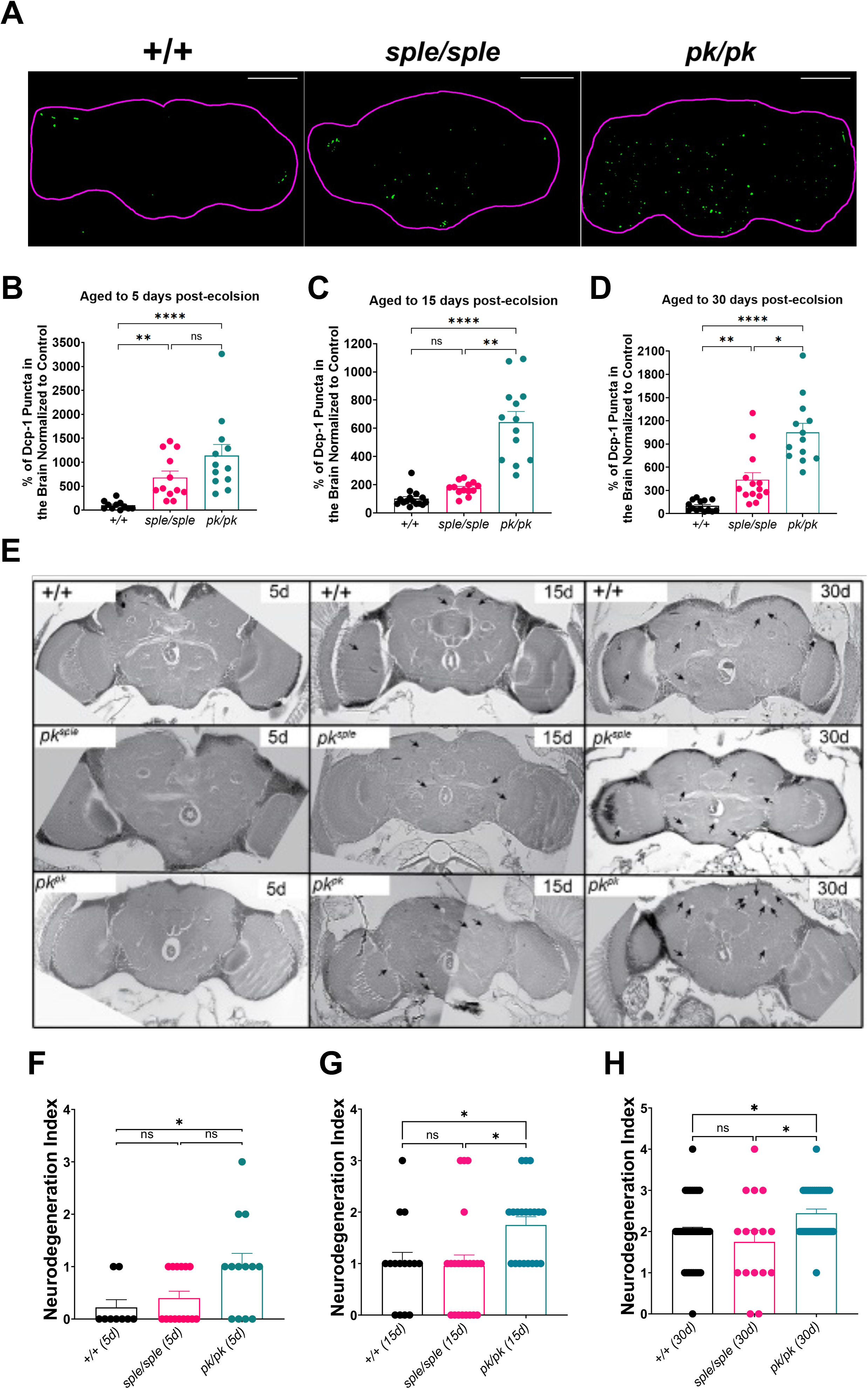
Homozygous *pk^pk^*mutants show significant increase in neurodegeneration across adulthood. A, Representative confocal images of 15-days old +/+, *sple/sple* and *pk/pk* mutant brains stained for Dcp-1 (green), a marker for apoptosis. Scale bar = 100 µm. **B-D,** Quantification of Dcp-1 positive puncta normalized to controls at 5-days (**B**), 15-days (**C**) and at 30-days (**D**) reveals a significant increase in cell death in both *sple/sple* and *pk/pk* mutant brains when compared to age matched controls. At 15-days and 30-days, *pk/pk* mutant brains also show a significant increase in cell death when compared to *sple/sple* mutant brains. Data shown are Mean ± SEM analyzed by Kruskal-Wallis test with multiple comparisons test. n = 11-14 brains for all genotypes and timepoints. **E,** Representative paraffin embedded 5 µm tissue sections at approximately mid-brain of +/+, *sple/sple* and *pk/pk* mutants aged to 5-days, 15-days and 30-days post eclosion and stained for H&E. Arrows indicate regions of vacuolization resulting from cell death in the brain. **F-H,** Quantification of neurodegeneration index in +/+, *sple/sple* and *pk/pk* mutant brains aged to 5-days (**F**), 15-days (**G**) and 30-days (**H**). *sple/sple* mutant brains show no significant increase in neurodegeneration index at any of the timepoints when compared to controls but *pk/pk* mutant brains show a significant increase in neurodegeneration index compared to controls at all timepoints analyzed. Data are Mean ± SEM analyzed by Kruskal-Wallis test with multiple comparisons. n = 9-41 brains for all genotypes and timepoints. For all data, *p ≤ 0.05, **p ≤ 0.01 and ****p ≤ 0.0001 and ns indicates no significant difference.

Both *pk^sple^* and *pk^pk^* mutant brains show evidence of increased oxidative stress, a known DNA mutagen (here and in (Nukala et al., 2023)), and other fly neurological disease mutants are often associated with increased DNA damage (Pessina et al., 2021; Wang et al., 2021) and reduced lifespan (Katzenberger et al., 2015; Maitra et al., 2019; Petersen et al., 2012). We first assessed DNA damage in the brains of *pk^sple^* and *pk^pk^*mutants by staining 15 dpe brains with anti-H2AV, an antibody used to detect DNA double strand breaks (DSBs) (Madigan et al., 2002), and anti-Elav, a neuronal marker, and quantified the H2AV-positive cells. Interestingly, we observed a significant increase in the percentage of H2AV positive cells in *pk^pk^*mutant brains but not in *pk^sple^* mutant brains when compared to controls, suggesting that neurons in *pk^pk^* mutant brains are undergoing significantly higher DNA damage than in *pk^sple^* mutant brains (Figure 4A-C). Next, we performed a longevity assay to determine the median lifespan of *pk^sple^*and *pk^pk^* mutants. We found that both mutants showed a significant reduction in median lifespan (38 days, *p* < 0.0001, and 30 days, *p* < 0.0001, respectively) when compared to control (44 days), with *pk^pk^*mutants having a more pronounced reduction than *pk^sple^* mutants (*p* < 0.0037) (Figure 4D). Since increased *AMP* gene expression in the glia has been linked to reduced lifespan in flies (Kounatidis et al., 2017), we assessed whether knocking down the IMD arm of the glial innate immune response in *pk^pk^* mutants extended their lifespan. Indeed, although *pk^pk^*/*pk^pk^*; UAS-*RelRNAi*/+ improved lifespan (likely due to leaky expression of the UAS-*RelRNAi*), glial-specific knockdown of *Relish* (an NF-κB transcription factor in the IMD pathway) in *pk^pk^* mutant brains using the Galactose responsive transcription factor 4/Upstream Activation Sequence (Gal4/UAS) expression system (Brand & Perrimon, 1993) restored lifespan to control levels (Figure 4E). Taken together, these data suggest that the higher amounts of neurodegeneration and DNA damage in the brains of *pk^pk^* mutants (compared to those of *pk^sple^* mutants), in combination with the heightened innate immune response, is likely contributing to their significantly reduced lifespan.

**Figure 4.**
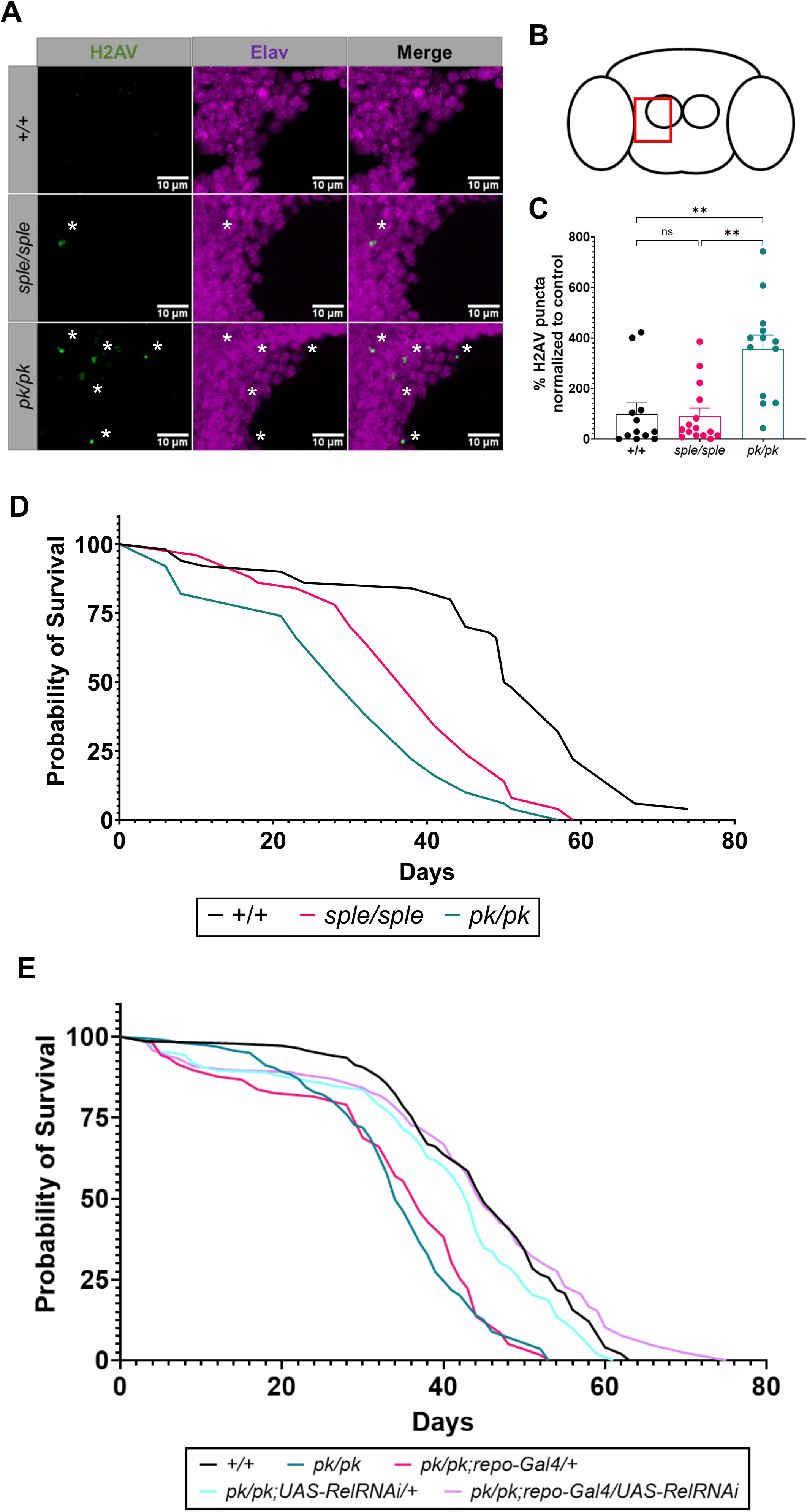
Homozygous *pk^pk^* mutants show significant DNA damage and reduced lifespan. A, Representative confocal images of 15-day old +/+, *sple/sple* and *pk/pk* mutant brains stained for anti-H2AV (green) and anti-Elav (magenta), a DNA damage marker and a neuron marker respectively. White asterisks indicate regions of co-localization between anti-H2AV and anti-Elav. Scale bar = 10 µm. **B,** shows the region of the brain (highlighted by the red box) represented in confocal images in **A. C,** Quantification of percentage of H2AV puncta normalized to control in 15-day old +/+, *sple/sple* and *pk/pk* mutant brains. There is a significant increase in H2AV staining in *pk/pk* mutant brains when compared to +/+ and *sple/sple* mutant brains. Data are Mean ± SEM analyzed by Kruskal-Wallis test with multiple comparisons test. N = 12-15 brains for all genotypes. **D,** Lifespan analysis of +/+, *sple/sple* and *pk/pk* mutants. Both *sple/sple* (38 days) and *pk/pk* (30 days) mutants show a significantly shorter lifespan than +/+ (44 days) with *pk/pk* mutants having a significantly shorter median lifespan than *sple/sple* mutants. Data are analyzed by Log-rank (Mantel-Cox) test. n = 48-50 flies for all genotypes. **E,** Lifespan analysis of +/+, *pk/pk*, *pk/pk*;*repo-Gal4/+, pk/pk;UAS-RelRNAi/+* and *pk/pk;repo-Gal4/UAS-RelRNAi* flies. The survival curve analysis shows a significant increase in median lifespan of *pk/pk;repo-Gal4/UAS-RelRNAi* (45 days) when compared to *pk/pk* mutants (34 days). Data are analyzed by Log-rank (Mantel-Cox) test. n = 72-111 flies for all genotypes. For all data, **p ≤ 0.01 and ns indicates no significant difference.

### 3.4. *pk^sple^* but not *pk^pk^* mutants show significant motor dysfunction despite lower levels of neuronal cell death

We previously reported that homozygous *pk^sple^* but not *pk^pk^* mutants are seizure-prone (Ehaideb et al., 2016). To assess whether this correlation extended across adulthood, we analyzed both *pk^sple^*and *pk^pk^* mutants compared to controls for seizure activity at 5, 15, and 30 dpe. For both males and females across all time points, only *pk^sple^* but not *pk^pk^* mutants showed evidence of seizure activity (Supplementary Figure 5A-B). However, given that both increased innate immune response and neuronal cell death in other fly models of neurological diseases have been linked with locomotor defects (Katzenberger et al., 2013; Kounatidis et al., 2017; Petersen et al., 2012), we tested whether homozygous *pk^pk^* mutants showed evidence of climbing defects or ataxia with age. While *pk^pk^* mutant males showed no significant decrease in climbing pass rate at any time point (5, 15, 30 dpe; Figure 5A) in contrast to *pk^sple^* mutant males, *pk^pk^* mutant females did show significantly decreased pass rates, although they were not as pronounced as for *pk^sple^* mutant females (Figure 5B). Next, we utilized pySolo (Gilestro & Cirelli, 2009) to determine whether *pk^pk^*and *pk^sple^* mutants showed evidence of ataxia across adulthood. Contrastingly, seizure-prone *pk^sple^*but not *pk^pk^* mutants showed significant increases in median turning angle (a measure of ataxia) across all time points when compared to age-matched controls (Figure 5C). Quantification of total distance traveled (pixels; evidence of compromised motor function including ataxia) revealed that both homozygous *pk^pk^*and *pk^sple^* mutants show a significant decrease in distance at 5 and 15 dpe while only *pk^pk^*mutants show a significant decrease at 30 dpe when compared to controls (Figure 5D). Collectively, these data suggest that despite the significantly higher neuronal cell death/neurodegeneration and reduced lifespan compared to seizure-prone *pk^sple^* mutants, *pk^pk^* adults only show modest motor deficits across adulthood.

**Figure 5.**
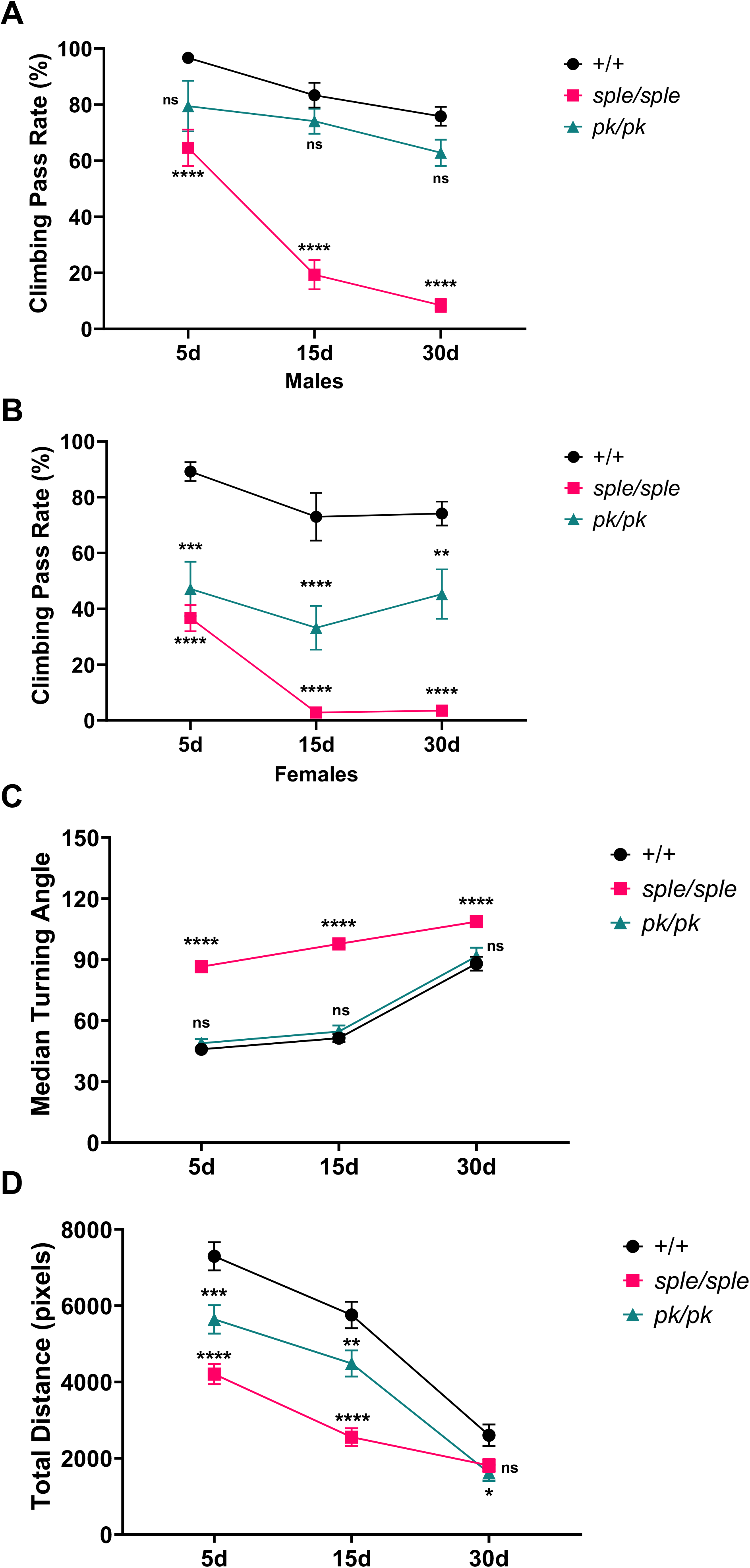
Homozygous *pk^sple^*but not *pk^pk^* mutants display increased locomotor defects with age. A, Quantification of percentage of climbing pass rate of 5, 15 and 30 day old male +/+, *sple/sple* and *pk/pk* mutants. Male *sple/sple* mutants show significant decrease while there is no significant difference in the percentage of climbing pass rate of *pk/pk* mutants when compared to controls across all timepoints. Data are Mean ± SEM analyzed by a Two-way ANOVA with Tukey’s multiple comparisons test. n = 4-10 replicates for all genotypes and timepoints. **B,** Quantification of percentage of climbing pass rate of 5, 15 and 30 day old female +/+, *sple/sple* and *pk/pk* mutants. Both female *sple/sple* and *pk/pk* mutants show significant decrease in the percentage of when compared to controls across all timepoints. Data are Mean ± SEM analyzed by a Two-way ANOVA with Tukey’s multiple comparisons test. n = 7-10 replicates for all genotypes and timepoints. **C,** Quantification of median turning angle in 5, 15 and 30 day old +/+ *sple/sple* and *pk/pk* mutants using pySolo. *sple/sple* mutants but not *pk/pk* mutants show a significant increase in median turning angle across all timepoints when compared to controls. Data are Mean ± SEM analyzed by Two-way ANOVA with Tukey’s multiple comparisons test. n = 57-100 flies for all genotypes and timepoints. **D,** Quantification of Total distance (pixels) in 5, 15 and 30 day old +/+ *sple/sple* and *pk/pk* mutants using pySolo. Both *sple/sple* mutants and *pk/pk* mutants show a significant decrease in total distance traveled when compared to controls with *sple/sple* mutants showing a greater decrease than *pk/pk* mutants when compared to controls at 5 and 15 days. At 30 days, *pk/pk* but not *sple/sple* mutants show a significant decrease in total distance (pixels) when compared to controls. Data are Mean ± SEM analyzed by Two-way ANOVA with Tukey’s multiple comparisons test. n = 60-100 flies for all genotypes and timepoints. For all data, * p ≤ 0.05, **p ≤ 0.01, ***p ≤ 0.001 and ****p ≤ 0.0001 and ns indicates no significant difference.

### 3.5. Seizure-prone *pk^sple^* mutants have pronounced learning and memory defects

Given that human *PRICKLE* patients as well as mouse mutants are associated with ASD-like learning and behavioral deficits (Paemka et al., 2013; Sowers et al., 2013), and that *PRICKLE* damaging variants are enriched in ASD patients (Paemka et al., 2013; Sowers et al., 2013), we wanted to determine whether homozygous *pk^sple^* and *pk^pk^*mutants showed impairments in learning and memory. First, we set out to assess whether *pk^sple^* and *pk^pk^* mutants were able to perceive smell. Upon exposure to increasing concentrations of 3-octanol (which flies will begin to find aversive) (Wang et al., 2003) and a standard concentration (1:500) of 4-methylcyclohexanol, we observed that both *pk^sple^*and *pk^pk^* mutants were able to perceive and avoid the increasing concentration of 3-octanol at similar levels when compared to controls suggesting that both homozygous *pk^sple^* and *pk^pk^* mutants are not affected by their ability to smell (Supplementary Figure 6). Next, we determined whether *pk^sple^* and *pk^pk^* mutants were able to effectively sense and respond to electric shocks. To do so, we performed shock avoidance assays using the T-maze apparatus originally developed by Tully and Quinn (Tully & Quinn, 1985) and quantified the results using the performance index (P.I.; 1.0 = perfect avoidance, 0.0 = completely random distribution). We found that while male *prickle* mutants (both *pk^sple^* and *pk^pk^*) showed no significant defects in responding to electric shocks (Figure 6A), *pk^sple^* mutant females do show a reduced ability to respond to the electric shocks (0.611 P.I.) relative to controls (0.782 P.I.) (Figure 6B). However, as the *pk^sple^* females still exhibit a strong P.I. above the 0.6 threshold, we reasoned that they should still be able to respond to aversive olfactory conditioning. Using an aversive olfactory conditioning assay, we thus tested the ability of the male and female *pk^sple^* and *pk^pk^* mutants to correctly associate an electric shock with different olfactory cues. We found that, relative to control males (0.659 P.I.), there was a statistically significant learning deficit in *pk^sple^* mutant males, but not *pk^pk^* males (0.329 and 0.600 P.I., respectively) (Figure 6C). Like *pk^sple^* mutant males, *pk^sple^* females (0.298 P.I.) also exhibit a statistically significant reduction in learning relative to controls (0.6918 P.I.), while *pk^pk^* mutant females are not significantly different from controls (0.656 P.I.) (Figure 6D). Intriguingly, while there is no significant difference in learning between *pk^pk^*mutants and controls, both male and female seizure-prone *pk^sple^*mutants learn significantly worse than controls, thus correlating ASD-associated learning deficits with seizure incidence.

**Figure 6.**
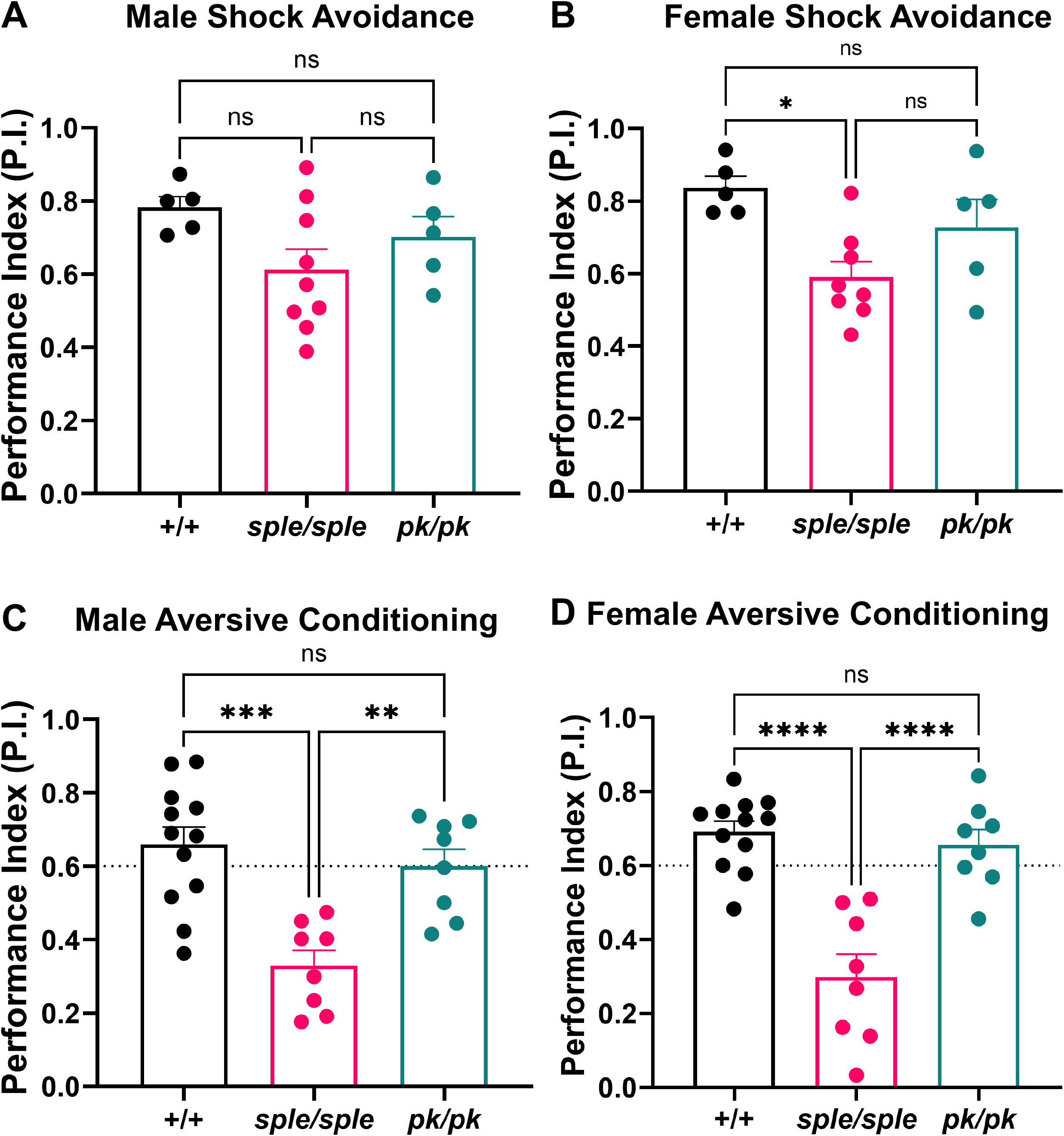
Homozygous *pk^sple^* mutants exhibit significant deficits in response to olfactory conditioning. A-B, Performance index (P.I.) of control, *pk/pk* and *sple/sple* mutants demonstrating avoidance of electric shocks in A) males and B) females. *sple*/*sple* mutant females, but not males, exhibit a statistically significant deficit relative to controls in their ability to perceive and respond to electric shocks, while *pk/pk* mutants show no difference relative to controls. **C-D,** P.I. of control, *pk/pk* and *sple/sple* mutants that demonstrate the ability to correctly associate an electric shock with a previously unconditioned scent. *sple/sple* mutants, but not *pk/pk* mutants exhibit statistically significant learning deficits relative to controls. n = 5-12 biological replicates with 50 flies per replicate. All data are reported as Mean ± SEM with ordinary one-way ANOVA with Tukey’s multiple comparisons test; for all data, * p ≤ 0.05, ** p ≤ 0.01, ***p ≤ 0.001, ****p ≤ 0.0001, and ns indicates no significant difference.

### 3.6. *pk^sple^* but not *pk^pk^* mutants demonstrate increased sensitivity to pain

Given the strong correlation between seizure activity and learning deficits in the *pk^sple^* mutant, we decided to assess whether additional features found in individuals with ASD might also be observed in *pk^sple^*flies. One such feature is an increase in pain sensitivity (Failla et al., 2020; Hoffman et al., 2023; Qian et al., 2024). To determine whether homozygous *pk^pk^* and *pk^sple^* mutants show any evidence of altered sensitivity to noxious heat, we performed a thermal nociception assay where we subjected 7–9 days old male controls as well as *dTrpA1^ins^* (a positive control which has known defects in heat perception) (Neely et al., 2011), *pk^sple^*, and *pk^pk^* mutants to increased temperatures ramping from 25°C to 50°C. We then recorded their nociceptive (jump) responses (Figure 7A) and tracked their velocity (Figure 7B). We observed that controls flies show a stereotypical jump profile on the ramped hotplate assay (Massingham et al., 2021). However, while homozygous *pk^pk^* mutants display a significant decrease in jumps at all temperatures similar to the known heat insensitive mutant *dTrpA1^ins^* (Neely et al., 2011), suggesting *pk^pk^* mutants have defects in sensing or escaping from noxious heat (Figure 7C), *pk^sple^* mutants display an increase in their nociceptive (jump) responses to lower temperatures (beginning at 40°C and peaking at 42°C; Figure 7A). Upon quantification of the velocities during the assay, we observed that homozygous *pk^pk^* mutants display an initially similar velocity to controls, suggesting no motor defects, while *pk^sple^* mutants have reduced initial velocities which could suggest a motor defect (Figure 7B, 7D). However, since *pk^sple^* mutants show a similar number of jump responses compared to control, this would further suggest that any potential motor defect does not impact the animal’s ability to activate the noxious jump escape response. Taken together, the thermal nociception assay revealed that homozygous *pk^pk^* mutants are insensitive to noxious heat pain while homozygous *pk^sple^* mutants have increased sensitivity to pain.

**Figure 7.**
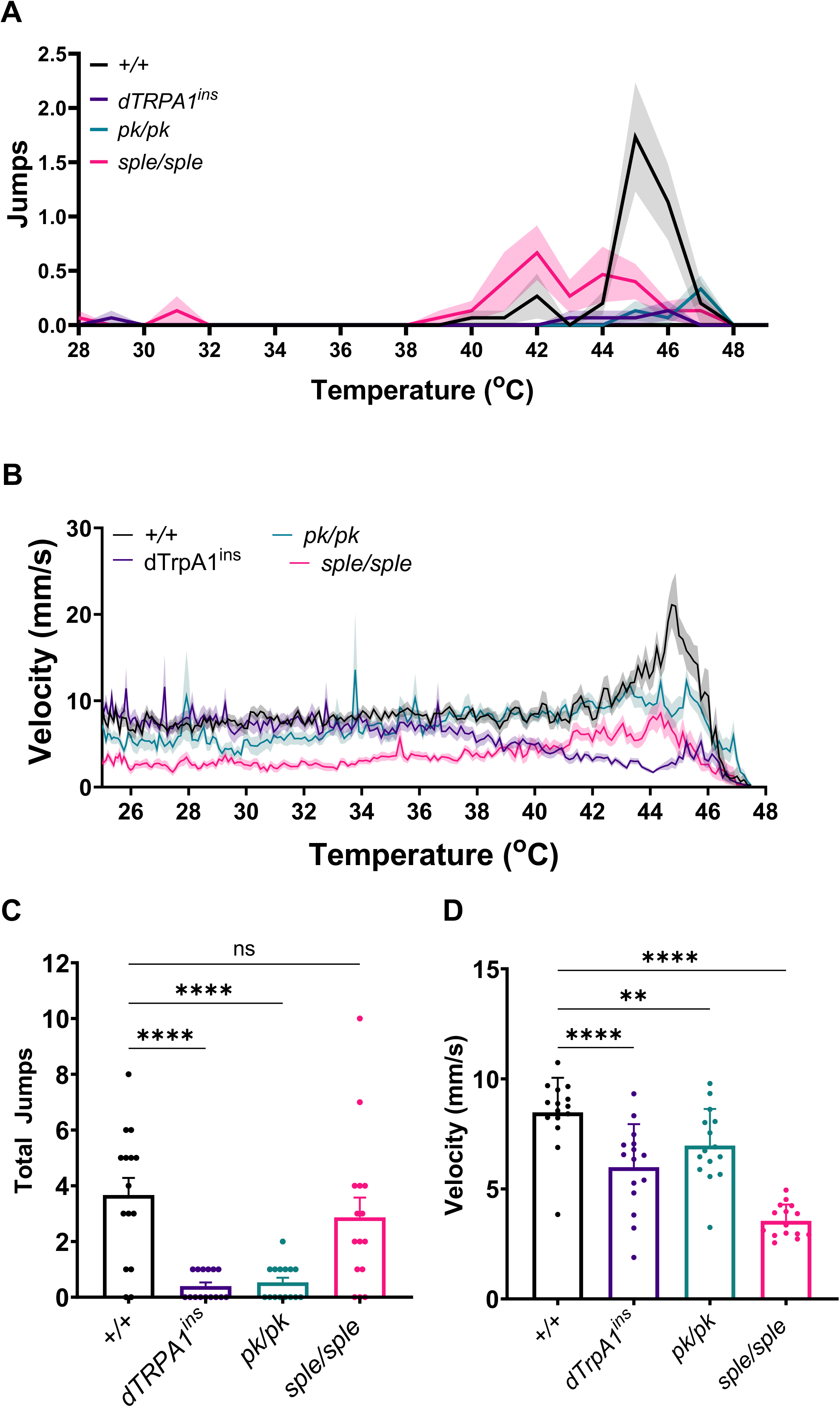
Homozygous *pk^sple^* and *pk^pk^* mutants have significant differences in nociceptive heat responses. A, Nociceptive (jump) responses to heat stimulation in control, *dTrpA1^ins^*, *pk/pk* and *sple/sple* mutants. **B,** Velocity (mm/s) binned in 1 second intervals in control, *dTrpA1^ins^*, *pk/pk* and *sple/sple* mutants. **C,** Total Jumps, (F_(4,65)_ = 9.765, *p* < 0.0001) quantified in control, *dTrpA1^ins^*, *pk/pk* and *sple/sple* mutants. **D,** Mean Velocity (mm/s) (F_(4,66)_ = 28.73, *p* < 0.0001) quantified in control, *dTrpA1^ins^*, *pk/pk* and *sple/sple* mutants. Statistically significant results are indicated on the graph; **p ≤ 0.01, ***p ≤ 0.001, ****p ≤ 0.0001; Ordinary One-way ANOVA with a Holm-Sidak correction for multiple comparisons. All data are reported as Mean ± SEM; n = 15 flies for all genotypes.

### 3.7. *pk^sple^* and *pk^pk^* mutants experience circadian rhythm defects

ASD patients show a significant comorbidity with disrupted sleep and circadian rhythm patterns (Cohen et al., 2014; Pinato et al., 2019; Xavier, 2021). Therefore, to determine whether homozygous *pk^sple^* and *pk^pk^*mutants have similar disruptions, we monitored the activity of control, *pk^pk^*and *pk^sple^* mutants both during 12 h/12 h light/dark entrainment (LD) and constant dark conditions (DD). While under entrainment, control flies showed the expected anticipation activity, measured as an increase in locomotion immediately preceding both the morning and evening changes in lighting (Figure 8A, black and gray arrowheads, respectively). We observed that homozygous *pk^pk^* but not *pk^sple^* mutants showed a significant decrease in both morning and evening anticipation activity (Figure 8B-E). Together, these data suggest that disruptions to the *pk^pk^* but not *pk^sple^* isoform expression may contribute to the entrainment defects of circadian rhythm.

**Figure 8.**
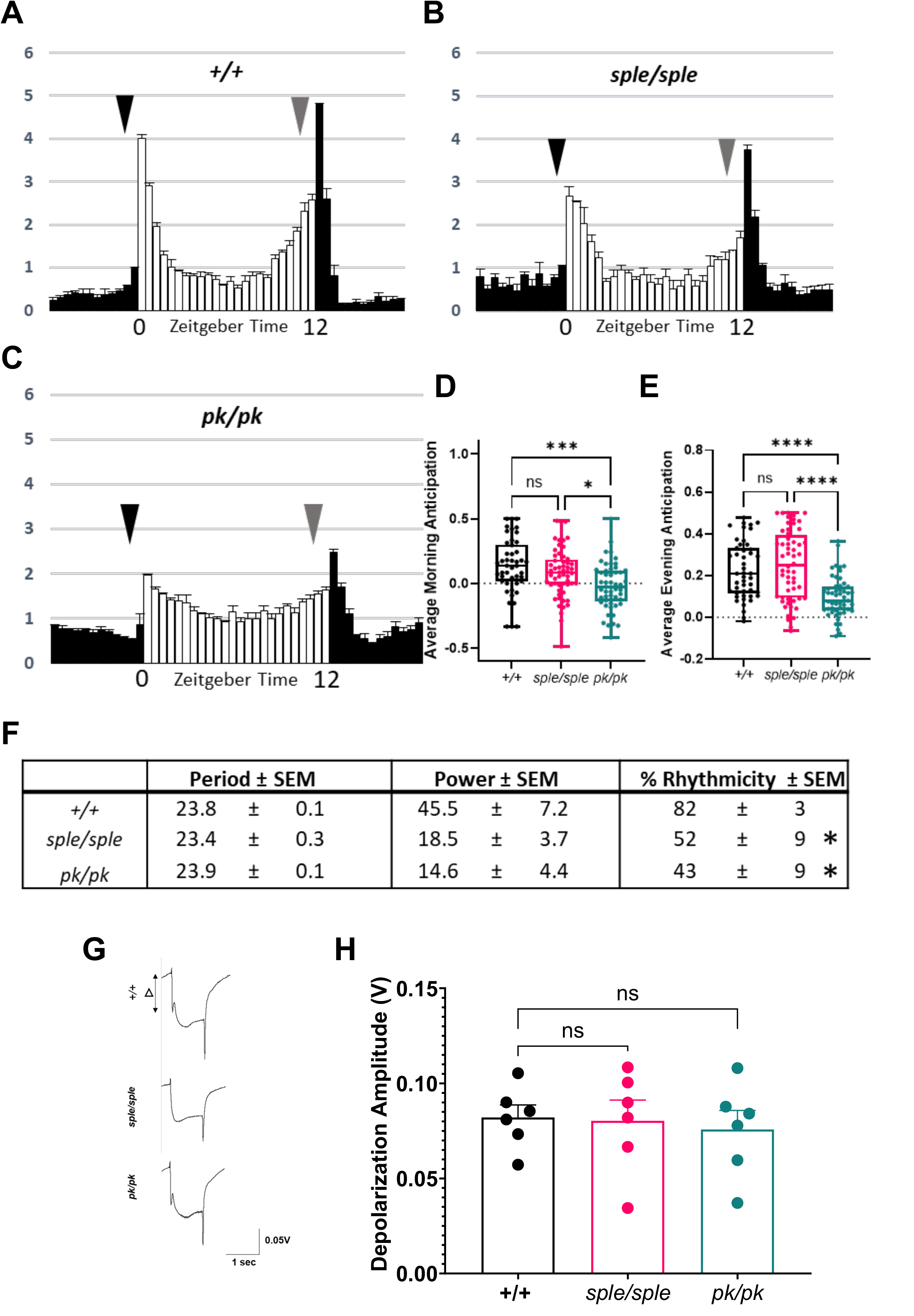
Homozygous *pk^sple^*and *pk^pk^* show circadian rhythm defects. A, Normalized locomotor activity patterns of control flies averaged over 4 days of 12h light:12h dark conditions (LD). Zeitgeber time indicated below the panel and white and black bars represent normalized activity in light and dark phase, respectively. Black and gray arrowheads indicate approximate timing of morning and evening anticipation respectively. n = 47 flies. **B,** Normalized locomotor activity patterns of *sple/sple* mutants averaged over 4 days LD. n = 56 flies. **C,** Normalized locomotor activity patterns of *pk/pk* mutants averaged over 4 days LD. n = 53 flies. **D-E,** Quantification of morning (**D**) and evening (**E**) anticipation in 12:12 LD conditions. *pk/pk* but not *sple/sple* mutants show a significant disruption to the circadian rhythm for both morning and evening anticipation behavior. Data shown are Box and Whisker plots (horizontal lines within boxes represent medians) with Kruskal Wallis test with post-hoc multiple comparisons test. **F,** Table shows Period and Power of rhythmicity along with percentage of rhythmicity in control, *sple/sple* and *pk/pk* mutant flies subjected to 7 days constant dark conditions (DD), as determined using chi-squared periodogram analysis. In both *sple/sple* and *pk/pk* mutants, % of rhythmic flies was significantly decreased when compared to controls. For flies showing rhythmicity, *pk/pk* mutants show a significant decrease in in power of rhythmicity while *sple/sple* mutants show a decreasing trend towards significance. (p = 0.106). In both *sple/sple* and *pk/pk* mutants, period length was not significantly disrupted when compared to controls. Statistical comparisons for DD data were made using Kruskal-Wallis test with post-hoc analysis. **G,** Representative ERG traces of 15-day old control, *sple/sple* and *pk/pk* mutants showing depolarization amplitudes. **H,** Quantification of depolarization amplitudes in control, *sple/sple* and *pk/pk* mutants shows no significant differences among the various genotypes. Data shown are Mean ± SEM with ordinary One-way ANOVA with post-hoc multiple comparisons test. For all data, *p ≤ 0.05, ***p ≤ 0.001, ****p ≤ 0.0001 and ns indicates no significance.

Conversely, under constant dark conditions, both *pk^pk^*and *pk^sple^* isoform expression in the fly were shown to be required for proper maintenance of circadian rhythm. We observed that while 82% of control flies were able to maintain rhythmicity in 7 days of DD conditions, only 43% of *pk^pk^* and 52% of *pk^sple^* mutants were able to maintain their rhythmicity under DD conditions. Moreover, for those *pk^pk^* and *pk^sple^* mutant flies that maintained the rhythmicity, they did so with demonstrably less power (a relative measurement of the strength of rhythmicity) when compared to controls (Figure 8F). Additionally, electroretinogram recordings of 15-day old control, *pk^sple^*and *pk^pk^* mutants revealed that there were no significant differences in depolarization potential among both *pk^pk^* and *pk^sple^* mutants when subjected to white light pulses, indicating that both the mutants are able to perceive light (Figure 8G-H). Collectively, these data suggest that both *pk^pk^*and *pk^sple^* isoforms are required to maintain circadian rhythm in flies, and that maintenance of the circadian rhythmicity is disrupted in both *prickle* mutants.

### 3.8. *pk^sple^* but not *pk^pk^* mutants exhibit increased social isolation relative to controls

ASD patients manifest difficulties in communication and social interactions (Fuentes et al., 2021), and social isolation assays have been used in flies to model such alterations (Hahn et al., 2013; Moscato et al., 2020; Stahl & Tomchik, 2024; Stojkovic et al., 2024; Wise et al., 2015). To assay whether *prickle* mutants demonstrate deficiencies in socialization, we assessed interindividual fly distances relative to nine conspecifics within a defined enclosed circular socialization chamber (SC) over the course of 15 minutes. As shown in Figure 9A, *pk^sple^* but not *pk^pk^* mutants exhibit statistically significant increases in social isolation 8-9 minutes after introduction into the SC compared to controls (average median interindividual distances 46.25 mm compared to 36.06 mm and 39.48, respectively). After 11-12 minutes in the SC, controls have now separated from one another to the same extent as the *pk^sple^* mutants (47.49 mm and 42.90 mm, respectively), whereas *pk^pk^* mutants are slower to explore the space (40.60 mm; Figure 9B). At 14-15 minutes (Figure 9C), the controls, *pk^sple^*, and *pk^pk^*mutants exhibit similar average median interindividual distances (44.75 mm, 48.09 mm, and 44.69 mm, respectively), demonstrating that all genotypes are now spread out across the SC. Overall, these data indicate that *pk^sple^*but not *pk^pk^* mutants socially isolate from their fly cohorts relative to controls, and remain socially isolated, consistent with difficulties in social interaction.

**Figure 9.**
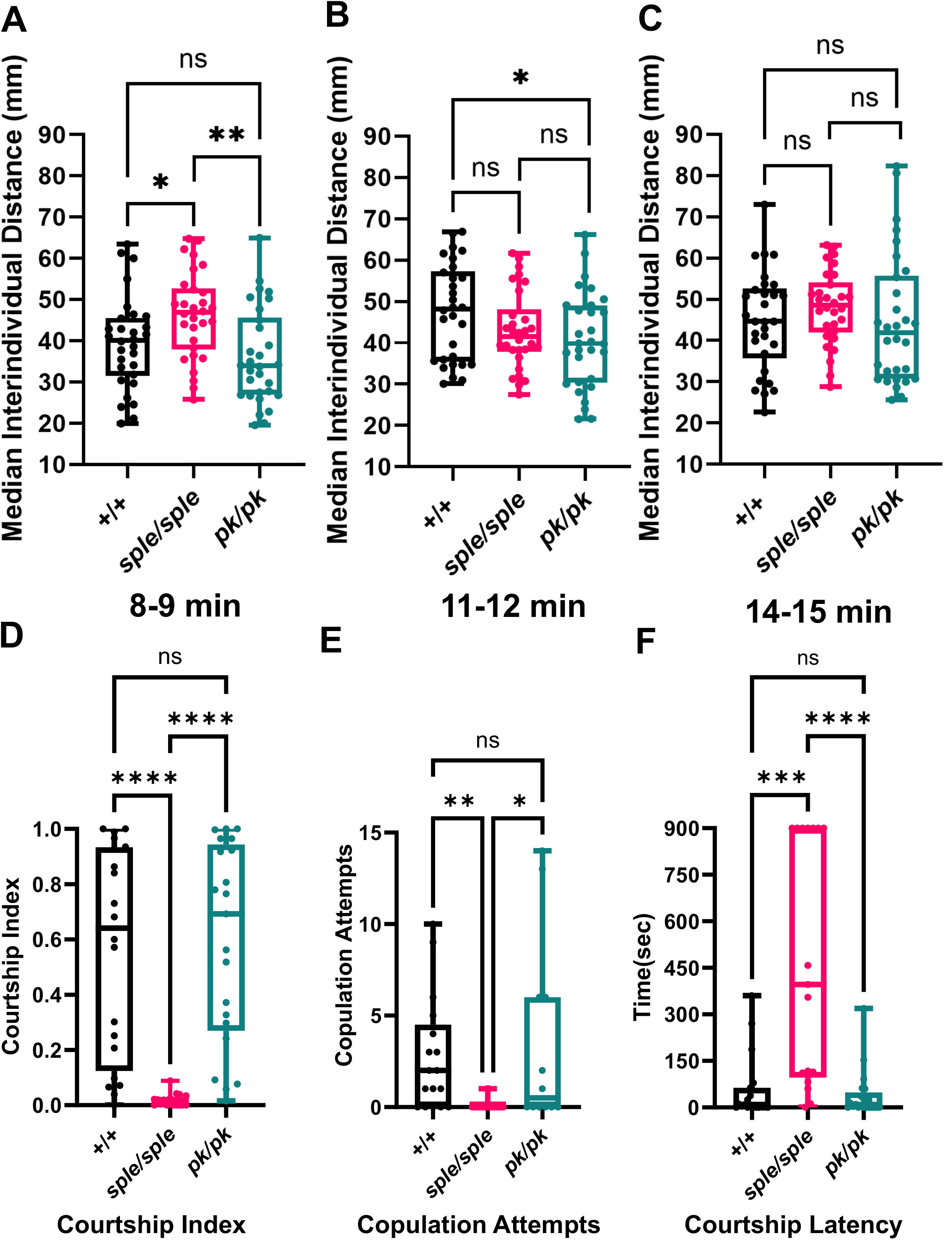
Homozygous *pk^sple^,* but not *pk^pk^* mutants, exhibit increased social isolation and reduced male courtship behaviors relative to controls. A-C, Median distances to nine conspecifics (in mm) for both controls and mutants at 8-9 min **(A)**, 11-12 min **(B)**, and 14-15 min **(C)** after transfer to the SC. *sple/sple* mutants show high levels of social isolation relative to both controls and *pk/pk* mutants soon after transfer to the SC and maintain increased social isolation throughout all timepoints assayed. Data represented as Box and Whisker plots (horizontal lines within boxes represent medians) with multiple Mann-Whitney tests, n = 10 flies per trial, 3 trials per genotype. **D-F,** Male *sple/sple* mutants show reduced courtship index **(D)**, increased courtship latency **(E)**, and reduced attempts to copulate **(F)** compared to male *pk/pk* mutants and controls. Data shown are Box and Whisker plots (horizontal lines within boxes represent medians) using the Kruskal-Wallis test, n ≥ 20 male flies tested for each genotype. For all data, *p ≤ 0.05, ** p ≤ 0.01, ***p ≤ 0.001, ****p ≤ 0.0001, and ns indicates no significance.

### 3.9. *pk^sple^* but not *pk^pk^* mutants show reduced male courtship behaviors

Courting is an essential social behavior in *Drosophila* and can be used as an assessment of social communication (Tener et al., 2024). To test whether *prickle* mutants have reduced courtship behaviors, we quantified naive *pk^sple^* and *pk^pk^* male courtship behavior towards virgin females in a circular chamber over 15 minutes. While *pk^sple^*male courtship behavior was reduced compared to controls (median courtship indices =.005, 0.640, respectively), *pk^pk^*mutant males showed similar courtship behavior to controls (median courtship index = 0.692) (Figure 9D).

Additionally, *pk^sple^* males showed a statistically significant increase in courtship latency (i.e., length of time before first courtship behavior; median = 396 sec) compared to controls and *pk^pk^*males (median = 8.8 and 6.0 sec, respectively) (Figure 9E). Finally, *pk^sple^*males showed a significant decrease in copulation attempts (median 0) whereas *pk^pk^* males showed similar copulation attempts to controls (medians 0.5 and 2, respectively) (Figure 9F). In summary, these data demonstrate that only *pk^sple^*males show significant reductions in the ability to court females, indicating that social communication is compromised in *pk^sple^* mutant flies.

### 3.10. *pk^sple^* but not *pk^pk^* mutant flies exhibit increased grooming behaviors

A core feature of ASD individuals is restrictive or repetitive behaviors (Caldwell-Harris, 2021). To determine whether our *prickle* mutant flies showed evidence of repetitive behaviors, we assessed whether *pk^pk^*and *pk^sple^* mutant flies had increased grooming behaviors (a characteristic of other fly ASD models) compared to controls (Hope et al., 2019; Marcogliese et al., 2022; Song et al., 2024). As shown in Figure 10A, *pk^sple^* mutant males exhibit more grooming bouts (21.0) than either control or *pk^pk^* males (8.7 and 8.8 bouts per video, respectively). Notably, *pk^pk^*mutant males exhibit no statistically significant difference in mean grooming bout number relative to controls. Bout duration is also isoform loss-dependent, as the mean bout durations in *pk^sple^* mutant males (106.6 frames per bout) last significantly longer than either control or *pk^pk^*males (51.95 and 53.25 frames per bout, respectively) (Figure 10B), with *pk^pk^*mutant males not significantly differing from controls. The increased bout number and duration observed in *pk^sple^*mutant males thereby results in *pk^sple^* flies (28.95%) spending a significantly higher percentage of time grooming than controls or *pk^pk^*mutant males (5.54% and 5.95%, respectively) (Figure 10C). As with bout number and duration, there was no statistically significant difference in percentage of time grooming between controls and *pk^pk^* mutant males. The same trends are observed in female flies. *pk^sple^* mutant females exhibit an increased mean number of grooming bouts (29.9 bouts per video) relative to controls and *pk^pk^* mutants (11.8 and 10.6 bouts per video, respectively) (Figure 10D) while *pk^pk^* mutant females do not exhibit a statistically significant difference in grooming bout number compared to controls. The grooming bouts in *pk^sple^*mutant females last significantly longer (107.40 frames per bout) than controls or *pk^pk^* mutant females (63.15 and 60.30 frames per bout, respectively) (Figure 10E) and the bout duration of the controls and *pk^pk^*mutant females was not significantly different. Similar to the males, *pk^sple^*mutant females spend a significantly larger percent of time grooming (55.13%) than both controls and *pk^pk^* mutants (8.93 and 8.29%, respectively) (Figure 10F). Taken together, these data indicate that *pk^sple^*mutant flies, but not *pk^pk^* mutants, exhibit restrictive behaviors which are a core feature of individuals with ASD.

**Figure 10.**
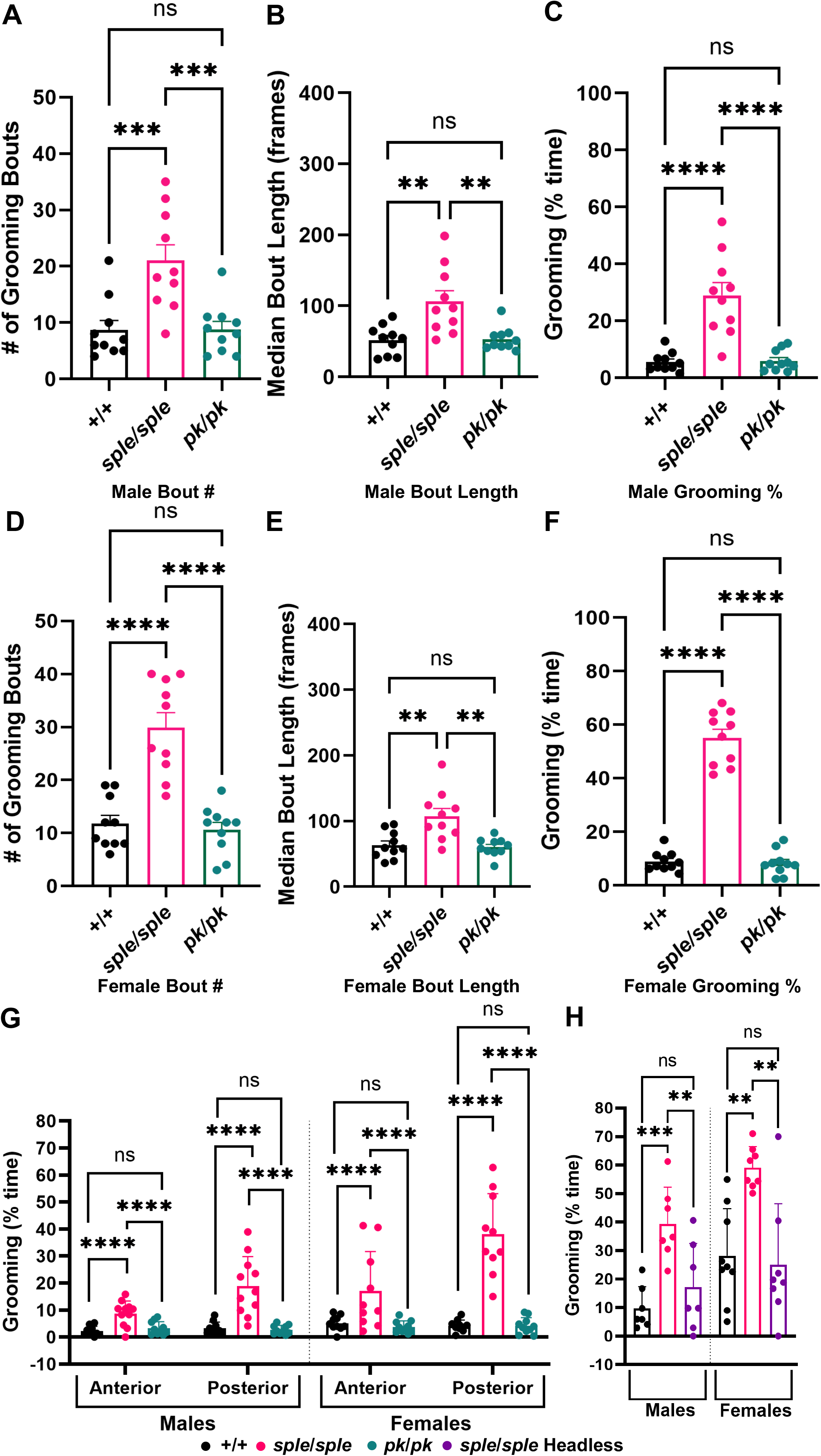
Homozygous *pk^sple^*mutants exhibit a significant increase in grooming behavior compared to controls. A-C, Grooming behavior in male control, *sple/sple* and *pk/pk* mutants quantified by bout number **(A)**, median bout duration **(B)**, and percent time spent grooming **(C)** show a significant increase in *sple/sple* mutants when compared to both control and *pk/pk* mutants. **D-F,** Grooming behavior in female control, *sple/sple* and *pk/pk* mutants quantified by bout number **(D)**, median bout duration **(E)**, and percent time spent grooming **(F)** show a significant increase in *sple/sple* mutants when compared to both control and *pk/pk* mutants. **G**, Quantification of percent time spent grooming the anterior and posterior modules of the body separated by sex in control, *sple/sple* and *pk/pk* mutants shows a significant increase in *sple/sple* mutants when compared to both control and *pk/pk* mutants. **H,** Quantification of percent time spent grooming in head-intact control and *sple/sple* mutants and decapitated *sple/sple* mutants. Both male and female head-intact *sple/sple* mutants show a significant increase in percent time spent grooming when compared to head-intact control and decapitated *sple/sple* mutants. All data are reported as Mean ± SEM with ordinary one-way ANOVA with Tukey’s multiple comparisons test. n = 7-10 flies per genotype. For all data, ** = p ≤ 0.01, *** = p ≤ 0.001, **** = p ≤ 0.0001, and ns indicates no significance.

We next wanted to determine whether grooming was unique to a particular region of the body, which could indicate whether a particular grooming module (anterior or posterior) is preferentially engaged in the *prickle* mutants. To quantify this, we classified the grooming bouts by which pair of legs (front vs hind) were involved. As shown in Figure 10G, *pk^sple^* mutant males exhibit a significantly higher percentage of time spent grooming in the anterior region (8.73%) relative to both controls and *pk^pk^* mutants (2.25 and 3.25 %, respectively). The same trend is observed in the posterior region, with *pk^sple^*mutant males exhibiting a significantly higher percentage of time grooming the posterior region (19.35%) than either control or *pk^pk^* mutant males (3.29 and 2.71%, respectively). *pk^sple^* females also demonstrate an increase in grooming percentage in the anterior (17.04%) relative to controls and *pk^pk^* mutants (5.10 and 3.77%, respectively) and the same trend is observed in the posterior region, with grooming in *pk^sple^*mutant females (38.07%) significantly higher than both controls and *pk^pk^*females (4.21 and 4.20%, respectively). These data demonstrate that the increase in grooming behavior is not confined to one region of the body, as both *pk^sple^* males and females exhibit statistically significant increases in grooming for both the anterior and posterior modules. Taken as a whole, these data demonstrate that only the *pk^sple^* but not *pk^pk^* mutants exhibit robust repetitive behaviors, a core characteristic of ASD.

To determine whether the brain contributes to repetitive grooming behavior in *pk^sple^* mutants, grooming was assessed in decapitated *pk^sple^* mutants (Figure 10H). Decapitated flies can survive for hours until they die of either starvation or desiccation (King et al., 2020) and have been shown to exhibit both induced and spontaneous grooming behaviors (Yellman et al., 1997) as well as accurate kicking responses in order to remove mites from their wings (Li et al., 2016). Relative to head-intact *pk^sple^* mutant males (median 34.69%), decapitated *pk^sple^*males exhibited a statistically significant decrease in percent time spent grooming (median 9.77%). The same trend is observed in females, with decapitated *pk^sple^* mutants exhibiting a significant decrease (median 19.26%) compared to head-intact *pk^sple^* mutants (median 58.06%). Intriguingly, no statistically significant difference is observed between decapitated *pk^sple^* flies (medians 9.77% for males, 19.26% for females) and age-matched, head-intact control flies (medians 5.83% for males, 24.24% for females), demonstrating that neuronal circuits emanating from the brain drive the excessive grooming in *pk^sple^* mutant flies.

## 4. Discussion

While many genes have been linked to autism spectrum disorder (Eyring & Geschwind, 2021; Qiu et al., 2022; Wang et al., 2025), only a handful have also been associated with its frequent comorbidity, epilepsy (Viscidi et al., 2013), and these include *TSC1/2, FMR1, MECP2*, and *SHANK3* (Zahra et al., 2022). In this study, we provide further evidence that *PRICKLE* fulfills the qualifications for such a candidate. While our previous work showed that pathogenic variants of the human epilepsy genes *PRICKLE1* and *PRICKLE2* are enriched in individuals with ASD, and that mice carrying *Prickle1* and *Prickle2* gene disruptions exhibit ASD-like behaviors (e.g., altered social interactions, learning and activity abnormalities, and behavioral inflexibility (Paemka et al., 2013; Sowers et al., 2013)), the current study appreciably extends these findings using a seizure-prone fly *prickle* mutant (*pk^sple^*). We previously showed that numerous characteristics of the human *PRICKLE* epilepsy-ataxia syndrome are conserved in *pk^sple^* mutant flies including early-onset myoclonic/tonic-clonic seizures that worsen with age, along with ataxia (Ehaideb et al., 2016; Nukala et al., 2023), and the observation that ASD-associated clinical features are observed in both humans and flies carrying pathogenic *PRICKLE* variants underscores a remarkable conservation of *PRICKLE* function from flies to humans. Particularly notable is the association of learning deficits, increased pain sensitivity, social interaction and communication abnormalities, and repetitive behaviors in the *pk^sple^* but *not pk^pk^* mutant thereby correlating this constellation of ASD phenotypes to a *prickle* mutation specifically associated with seizures and thus providing evidence that these phenotypes are in fact genetically linked. Perhaps even more remarkable, the *pk^pk^* mutants have substantially more neuronal cell death as well as neurodegeneration (i.e., vacuolization) than the *pk^sple^*mutants in spite of the fact that they are largely devoid of ASD characteristics.

Processes such as associative learning, grooming, and social interaction clearly require higher level cognitive processing and are not simply controlled by autonomic, self-governing circuits. Behaviors such as these must be allocated to their appropriate time and context, as flies must balance a variety of behaviors including searching for food, avoiding threats, eating, flying, mating, etc. In contrast, *pk^sple^*flies obsessively and repeatedly groom their bodies such that all measures of grooming behavior (i.e., number of bouts, duration of bouts, location of grooming activity) are appreciably increased relative to controls (see Figure 10), and these behaviors are markedly reminiscent of the restrictive and repetitive behaviors in individuals with ASD, including body rocking, hand flapping, finger flicking, and pacing (Caldwell-Harris, 2021). Previous work in the field has demonstrated that central pattern generators (CPGs) play an important role in fly grooming (Ravbar et al., 2021), and recent work has shown that circuits emanating from the brain can influence not only whether a grooming bout is initiated, but where on the body that grooming is directed (i.e., the activation of positional body grooming modules)(Cande et al., 2018; Guo et al., 2022). The striking reduction in abnormal grooming behavior observed in headless *pk^sple^* (see Figure 10H) flies is consistent with brain circuitry controlling the grooming behavior, providing compelling evidence that grooming is not simply an autonomic CPG-generated behavior but is controlled by more complex processing. Additionally, communication and socialization among flies, which also require higher order cognitive processing, are compromised in *pk^sple^* flies. The courtship index (which quantifies the number of courtship communication behaviors of a male fly in the presence of a female fly) of *pk^sple^*males is close to zero, as are copulation attempts over the course of the 15 minute experimental time period, while time to first courtship behavior (courtship latency) is markedly increased relative to controls, thus demonstrating that *pk^sple^* mutant males are significantly compromised in all three parameters assessed. These observations are also reflected in the social isolation experiment whereby *pk^sple^* mutant flies fail to congress and interact with their fellow flies (a natural fly behavior) at any point in time measured, in contrast with control or *pk^pk^* flies which immediately interact after they are placed in the socialization chamber.

Learning capacity (significantly reduced in *pk^sple^* flies) is another activity requiring higher level cognitive processing, and up to 75% of individuals with ASD are intellectually disabled and exhibit learning deficits (Paemka et al., 2013). To assess learning, we used an aversive Pavlovian conditioning procedure that tests whether flies can associate an odor with an electric shock. It is important to point out that while reductions in the performance index (P.I.) could potentially arise from several reasons, such as an inability to perceive the odor or the pain from the electric shock, *pk^sple^* mutants were able to perceive and avoid an increasing concentration of 3-octanol similar to both control and *pk^pk^* mutants suggesting that olfaction is not impaired (Supplementary Figure 6). Furthermore, *pk^sple^* mutants have an increased sensitivity to pain (using the thermal nociception assay), which if anything should aid in their ability to perceive the electric shock, and the *pk^sple^*mutants perform well in a shock avoidance test, reaching high performance indexes of approximately 0.6 for both males and females (Figure 6), the generally accepted value for strong performance. Although *pk^sple^* females do show a modest reduction in shock avoidance compared to controls (P.I.∼0.6), these females show a much greater reduction of the performance index in the aversive conditioning assay (P.I.∼0.3), thereby demonstrating that their ability to learn is nonetheless compromised. The observation that defects in bifurcation of axons of the mushroom body (the learning and memory center of the fly brain) have been observed in *pk^sple^* but not *pk^pk^* mutants (which we have noted as well) provides one potential explanation for the *pk^sple^*-specific learning disability (Shimizu et al., 2011). We were, however, surprised to find that *pk^pk^* mutants performed so well in the learning assay given their extensive neuronal damage and death, thereby suggesting that the brain is a highly resilient organ when it comes to this form of associative learning.

The lack of pronounced motor deficits (i.e., seizure activity, ataxia, and climbing ability) in the *pk^pk^* mutants was not anticipated (Figure 3-4), since other fly neurological disease models showing neuronal cell death and neurodegeneration oftentimes show locomotor dysfunction (Katzenberger et al., 2013; Kounatidis et al., 2017; Maitra et al., 2019; Petersen et al., 2012). As we previously showed that the neuronal cell death leads to seizure exacerbation in the *pk^sple^* flies (Nukala et al., 2023), our data collectively suggest that different sets of neurons must be dying in the *pk^sple^* versus *pk^pk^* flies which in turn leads to the *pk^sple^*-specific seizure/ataxia phenotypes. As the neuronal cell death is widespread in both genotypes, it has been challenging to identify candidate neurons whose death leads to seizure activity, an area of research we are actively pursuing. Additionally, while both mutants are able to perceive light normally, both fail to maintain their rhythmicity under dark-dark conditions, but only the *pk^pk^* isoform is required for the entrainment to light-dark conditions thereby demonstrating a greater dependence on this isoform in circadian rhythm (Figure 8). Moreover, the *pk^pk^* mutants show a greatly reduced lifespan compared to *pk^sple^*, suggesting that the sustained neuronal damage eventually catches up to these flies. Interestingly, the lifespan reduction can be rescued by knocking down the glial IIR (Figure 4), underscoring the connection between nervous system health and longevity as has been previously observed (Kounatidis et al., 2017). Age-dependent increases in DNA damage as a result of decrease in activity of DNA repair processes has been well documented in flies (Pessina et al., 2021; Wang et al., 2021). In *pk^pk^*mutants, accelerated aging potentially due to a combination of aggravated IIR and oxidative stress could be contributing to the increased DNA damage in the brain, and previous studies have shown that oxidative stress is a key contributor to mitochondrial dysfunction, DNA damage and aging (Cui et al., 2012; Dai et al., 2014; Shadfar et al., 2023) while reduced antioxidant gene expression or excessive oxidative stress has been associated with various neurological diseases (Dias-Santagata et al., 2007; Galbusera et al., 2004; Huai & Zhang, 2019; Hussain et al., 2018; Niveditha et al., 2017).

Although the *pk^sple^* and *pk^pk^* mutations are loss-of-function and inactivate the *pk^sple^* and *pk^pk^* isoforms, respectively (Lilienthal et al., 2022), it is intriguing to note that the resulting neuronal phenotypes of the mutants, overall, are strikingly different especially given the known roles of these isoforms in the fly epidermis (see Supplementary Figure 7). There, the *prickle* isoforms help establish bristle and hair orientation across the adult body plan. Mutations in either isoform result in polarity defects that are cell-type specific with *pk^pk^* mutants showing altered polarity of the wing bristles, while *pk^sple^* mutants show altered polarity of leg bristles (Gubb et al., 1999). Across the epidermis, gradients of Dachsous (Ds), an atypical cadherin, and it’s interacting partner Dachs (D), are established such that some epidermal cells exhibit increased Ds/D at their proximal ends whereas others exhibit increased Ds/D distally (Sharp & Axelrod, 2016). These opposing gradient signals present a problem if seeking to employ them as a polarity-establishing code across an epidermal sheet. To remedy this issue, while Pk^pk^ localizes proximally in *pk^pk^*-expressing cells where Ds/D is highest at the distal end of the cell, the Pk^sple^ isoform (through its unique N-terminus) binds to Ds/D in *pk^sple^*-expressing cells in which the Ds/D is high at the proximal end thereby acting as the “gradient rectifier” (Ayukawa et al., 2014). The end result is that all cells localize one or the other Prickle isoform proximally, and once the relevant Prickle isoform is localized, it is thought that both isoforms carry out virtually identical cellular functions including organization of microtubule polarity (Sharp & Axelrod, 2016), perhaps not surprising given that, other than their N termini, the proteins are identical in structure (Supplementary Figure 1; (Gubb et al., 1999)). Our work here suggests that the rules within the nervous system are likely different. Neuronal architecture is quite distinct from that of epidermal sheets, whereby neuronal microtubules show plus-end out and minus-end out polarities in axons and dendrites, respectively (Stone et al., 2008), while both axons and dendrites respond to pathfinding/guidance cues in order to establish complex three-dimensional structures (Alfadil & Bradke, 2023). Second, the unique expression patterns of the *pk^pk^* and *pk^sple^* isoforms suggest a divergence from their roles in the epidermis.

*pk^sple^* and *pk^M^* are the most widely expressed isoforms in the crawling third instar larval brain (showing largely overlapping expression patterns), with *pk^pk^* showing much more localized expression (see Figure 1). Most of the cells expressing *pk^pk^*also express *pk^sple^*, and in a few isolated cases, some cells exclusively express *pk^pk^* including the neuroepithelial cells residing in the medial lamina of the optic lobe. Immediately distolateral to these cells is the lateral lamina which is positive for both *pk^pk^*and *pk^sple^*, cells which arose from the neuroepithelial cells, and moving even further distolaterally, these cells (which have now become neuroblasts and neurons) express only *pk^sple^*. Together, these data raise the intriguing possibility that *pk^pk^* labels the neuroepithelial fate whereby, once cells are committed to the neuroblast/neuron lineage, *pk^sple^* expression is activated and *pk^pk^* is turned off. Future studies assessing the expression patterns of these isoforms throughout larval development are needed to determine whether this hypothesis is correct. Nonetheless, it is remarkable that several of the phenotypes in the *pk^pk^* mutant are significantly more severe than in the *pk^sple^* mutant, including additional neuronal cell death, a more reduced lifespan, and neurodegeneration including vacuolization (which is not observed in *pk^sple^*mutants) in spite of the seemingly limited *pk^pk^* brain expression. Given the widespread prevalence of neuroblasts in the larval brain, if a neuroepithelial to neuroblast transition is indeed facilitated by *pk^pk^* expression, the loss of *pk^pk^* might have severe downstream consequences. In any case, these results imply that *pk^pk^* and *pk^sple^* may play at least partially independent roles in neurons. Consistent with this hypothesis, previous work in our laboratory has shown that *pk^sple^* mutants exhibit enhanced anterograde axon vesicle transport while *pk^pk^* mutants exhibit dramatically reduced axon vesicle transport, and that only overexpression of *pk^sple^* but not *pk^pk^* can reverse motor neuron axonal microtubule polarity (Ehaideb et al., 2014; Ehaideb et al., 2016).

Although the behavioral differences between *pk^pk^* and *pk^sple^* mutants are noteworthy, several pathophysiological similarities are observed at the cellular level. For example, both *pk^pk^*and *pk^sple^* mutant brains show a significant increase in IIR and oxidative stress response gene expression, although the oxidative stress response is somewhat more muted in the *pk^pk^* mutants with only 8 genes (mostly encoding for cytochrome p450s) being significantly upregulated while other key oxidative stress mitigator enzymes (dehydrogenases, catalases, reductases, transferases) are not (Figure 2). While neuronal cell death is more pronounced in the *pk^pk^* mutant, both *pk^pk^* and *pk^sple^* mutants do in fact show sustained, increased neuronal cell death across adulthood (Figure 3). Nonetheless, the fact that *pk^sple^* exhibits every ASD characteristic we assayed for, while *pk^pk^*exhibits only a single characteristic (i.e., circadian rhythm abnormalities), is worth noting as ASD is indeed a spectrum even amongst multiple affected family members, with some individuals displaying more severe clinical features than others (McDonald et al., 2020; Nisar et al., 2019). Comparing and contrasting the two *prickle* mutants has shown how different mutations in the same gene can have a greater or lesser impact on a neurodevelopmental disorder such as ASD depending on how severely the gene function is compromised. Taken as a whole, our data reveal that different mutations in the same gene can lead to a breadth of phenotypes, some of which may be similar but many of which are strikingly different thus underscoring the importance of broadscale analysis when studying genes associated with neurological diseases. Similar to other neurological disorders, a large percentage of epilepsies are accompanied by comorbid conditions that increase the phenotypic spectrum of the disease (Kanner, 2016), and only with careful assessment can these comorbidities be accurately quantified.

## CRediT authorship contribution statement

**Krishna M. Nukala:** Conceptualization, Investigation, Formal analysis, Methodology, Visualization, Writing-original draft, Writing-review and editing. **Brady Williquett:** Conceptualization, Investigation, Formal analysis, Methodology, Visualization, Writing-original draft, Writing-review and editing. **Anthony J. Lilienthal:** Investigation, Formal analysis, Visualization, Writing-original draft. **Dakota M. Thompson:** Investigation, Formal analysis. **Josephine N. Massingham:** Investigation, Formal analysis. **Shu Hui Lye:** Investigation, Formal analysis. **Avery Yu:** Investigation, Formal analysis. **Bridget C. Lear:** Methodology, Formal analysis, Writing-review and editing. **G. Gregory Neely:** Methodology, Formal analysis, Writing-review and editing. **Stanislava Chtarbanova:** Methodology, Formal analysis, Visualization, Writing-review and editing, Resources. **J. Robert Manak:** Conceptualization, Methodology, Writing-original draft, Writing-review and editing, Supervision, Project administration, Funding acquisition.

## Funding

This work was supported by grants National Institutes of Health grant R01NS098590 (JRM, Alexander G. Bassuk). SC was supported by start-up funds from the University of Alabama.

## Declaration of competing interests

The authors have no competing interests to declare.

## Acknowledgements

We thank Shannon Mahowald and Grant Salvucci for help with circadian rhythm and lifespan analysis, respectively, and Dr. Mrutyunjaya Parida for software implementation. We also thank Alex Bassuk for helpful and insightful discussions.

## Appendix A. Supplementary data

## Data availability

Relevant fly lines and data sets analyzed from the current study are available from the corresponding author upon reasonable request.

## Notes

### Competing Interest Statement

The authors have declared no competing interest.

## References

Alfadil, E., & Bradke, F. (2023). Moving through the crowd. Where are we at understanding physiological axon growth? Semin Cell Dev Biol, 140, 63–71. 10.1016/j.semcdb.2022.07.001

Ayukawa, T., Akiyama, M., Mummery-Widmer, J. L., Stoeger, T., Sasaki, J., Knoblich, J. A., Senoo, H., Sasaki, T., & Yamazaki, M. (2014). Dachsous-dependent asymmetric localization of spiny-legs determines planar cell polarity orientation in Drosophila. Cell Rep, 8(2), 610–621. 10.1016/j.celrep.2014.06.009

Ban, Y., Yu, T., Feng, B., Lorenz, C., Wang, X., Baker, C., & Zou, Y. (2021). Prickle promotes the formation and maintenance of glutamatergic synapses by stabilizing the intercellular planar cell polarity complex. Sci Adv, 7(41), eabh2974. 10.1126/sciadv.abh2974

Black, D. W., Grant, J. E., & American Psychiatric Association. (2014). DSM-5 guidebook : the essential companion to the Diagnostic and statistical manual of mental disorders, fifth edition (First edition. ed.). American Psychiatric Publishing.

Bolus, H., Crocker, K., Boekhoff-Falk, G., & Chtarbanova, S. (2020). Modeling Neurodegenerative Disorders in Drosophila melanogaster. Int J Mol Sci, 21(9). 10.3390/ijms21093055

Bonini, N. M., & Fortini, M. E. (2003). Human neurodegenerative disease modeling using Drosophila. Annu Rev Neurosci, 26, 627–656. 10.1146/annurev.neuro.26.041002.131425

Brand, A. H., & Perrimon, N. (1993). Targeted Gene-Expression as a Means of Altering Cell Fates and Generating Dominant Phenotypes. Development, 118(2), 401–415.

Brown, J. B., Boley, N., Eisman, R., May, G. E., Stoiber, M. H., Duff, M. O., Booth, B. W., Wen, J., Park, S., Suzuki, A. M., Wan, K. H., Yu, C., Zhang, D., Carlson, J. W., Cherbas, L., Eads, B. D., Miller, D., Mockaitis, K., Roberts, J.,…Celniker, S. E. (2014). Diversity and dynamics of the Drosophila transcriptome. Nature, 512(7515), 393–399. 10.1038/nature12962

Buescher, A. V., Cidav, Z., Knapp, M., & Mandell, D. S. (2014). Costs of autism spectrum disorders in the United Kingdom and the United States. JAMA Pediatr, 168(8), 721–728. 10.1001/jamapediatrics.2014.210

Caldwell-Harris, C. L. (2021). An Explanation for Repetitive Motor Behaviors in Autism: Facilitating Inventions via Trial-and-Error Discovery. Front Psychiatry, 12, 657774. 10.3389/fpsyt.2021.657774

Cande, J., Namiki, S., Qiu, J., Korff, W., Card, G. M., Shaevitz, J. W., Stern, D. L., & Berman, G. J. (2018). Optogenetic dissection of descending behavioral control in Drosophila. Elife, 7. 10.7554/eLife.34275

Cao, Y., Chtarbanova, S., Petersen, A. J., & Ganetzky, B. (2013). Dnr1 mutations cause neurodegeneration in Drosophila by activating the innate immune response in the brain. Proc Natl Acad Sci U S A, 110(19), E1752–1760. 10.1073/pnas.1306220110

Chien, Y. L., Wu, S. W., Chu, C. P., Hsieh, S. T., Chao, C. C., & Gau, S. S. (2017). Attenuated contact heat-evoked potentials associated with sensory and social-emotional symptoms in individuals with autism spectrum disorder. Sci Rep, 7, 36887. 10.1038/srep36887

Cho, B., Song, S., Wan, J. Y., & Axelrod, J. D. (2022). Prickle isoform participation in distinct polarization events in the Drosophila eye. PLoS One, 17(2), e0262328. 10.1371/journal.pone.0262328

Cohen, S., Conduit, R., Lockley, S. W., Rajaratnam, S. M., & Cornish, K. M. (2014). The relationship between sleep and behavior in autism spectrum disorder (ASD): a review. J Neurodev Disord, 6(1), 44. 10.1186/1866-1955-6-44

Cui, H., Kong, Y., & Zhang, H. (2012). Oxidative stress, mitochondrial dysfunction, and aging. J Signal Transduct, 2012, 646354. 10.1155/2012/646354

Dai, D. F., Chiao, Y. A., Marcinek, D. J., Szeto, H. H., & Rabinovitch, P. S. (2014). Mitochondrial oxidative stress in aging and healthspan. Longev Healthspan, 3, 6. 10.1186/2046-2395-3-6

DeSalvo, M. K., Mayer, N., Mayer, F., & Bainton, R. J. (2011). Physiologic and anatomic characterization of the brain surface glia barrier of Drosophila. Glia, 59(9), 1322–1340. 10.1002/glia.21147

Dey, S., Mondal, P., Mandal, S., Sasmal, S., Chakraborty, N., & Das, A. (2024). Paradigms for Behavioral Assessment in Drosophila Model of Autism Spectrum Disorder. J Vis Exp(211). 10.3791/66649

Dias-Santagata, D., Fulga, T. A., Duttaroy, A., & Feany, M. B. (2007). Oxidative stress mediates tau-induced neurodegeneration in Drosophila. J Clin Invest, 117(1), 236–245. 10.1172/JCI28769

Dorrego-Rivas, A., Ezan, J., Moreau, M. M., Poirault-Chassac, S., Aubailly, N., De Neve, J., Blanchard, C., Castets, F., Freal, A., Battefeld, A., Sans, N., & Montcouquiol, M. (2022). The core PCP protein Prickle2 regulates axon number and AIS maturation by binding to AnkG and modulating microtubule bundling. Sci Adv, 8(36), eabo6333. 10.1126/sciadv.abo6333

Dubois, A., Boudjarane, M., Le Fur-Bonnabesse, A., Dion, A., L’Heveder, G., Quinio, B., Walter, M., Marchand, S., & Bodere, C. (2020). Pain Modulation Mechanisms in ASD Adults. J Autism Dev Disord, 50(8), 2931–2940. 10.1007/s10803-019-04361-x

Egger, B., Boone, J. Q., Stevens, N. R., Brand, A. H., & Doe, C. Q. (2007). Regulation of spindle orientation and neural stem cell fate in the Drosophila optic lobe. Neural Dev, 2, 1. 10.1186/1749-8104-2-1

Ehaideb, S. N., Iyengar, A., Ueda, A., Iacobucci, G. J., Cranston, C., Bassuk, A. G., Gubb, D., Axelrod, J. D., Gunawardena, S., Wu, C. F., & Manak, J. R. (2014). prickle modulates microtubule polarity and axonal transport to ameliorate seizures in flies. Proc Natl Acad Sci U S A, 111(30), 11187–11192. 10.1073/pnas.1403357111

Ehaideb, S. N., Wignall, E. A., Kasuya, J., Evans, W. H., Iyengar, A., Koerselman, H. L., Lilienthal, A. J., Bassuk, A. G., Kitamoto, T., & Manak, J. R. (2016). Mutation of orthologous prickle genes causes a similar epilepsy syndrome in flies and humans. Ann Clin Transl Neurol, 3(9), 695–707. 10.1002/acn3.334

Ekengren, S., & Hultmark, D. (2001). A family of Turandot-related genes in the humoral stress response of Drosophila. Biochem Biophys Res Commun, 284(4), 998–1003. 10.1006/bbrc.2001.5067

Eyring, K. W., & Geschwind, D. H. (2021). Three decades of ASD genetics: building a foundation for neurobiological understanding and treatment. Hum Mol Genet, 30(20), R236–R244. 10.1093/hmg/ddab176

Fabisiak, T., & Patel, M. (2022). Crosstalk between neuroinflammation and oxidative stress in epilepsy. Front Cell Dev Biol, 10, 976953. 10.3389/fcell.2022.976953

Failla, M. D., Gerdes, M. B., Williams, Z. J., Moore, D. J., & Cascio, C. J. (2020). Increased pain sensitivity and pain-related anxiety in individuals with autism. Pain Rep, 5(6), e861. 10.1097/PR9.0000000000000861

Feany, M. B., & Bender, W. W. (2000). A Drosophila model of Parkinson’s disease. Nature, 404(6776), 394–398. 10.1038/35006074

Frundt, O., Grashorn, W., Schottle, D., Peiker, I., David, N., Engel, A. K., Forkmann, K., Wrobel, N., Munchau, A., & Bingel, U. (2017). Quantitative Sensory Testing in adults with Autism Spectrum Disorders. J Autism Dev Disord, 47(4), 1183–1192. 10.1007/s10803-017-3041-4

Fuentes, J., Hervas, A., & Howlin, P. (2021). ESCAP practice guidance for autism: a summary of evidence-based recommendations for diagnosis and treatment. Eur Child Adolesc Psychiatry, 30(6), 961–984. 10.1007/s00787-020-01587-4

Galbusera, C., Facheris, M., Magni, F., Galimberti, G., Sala, G., Tremolada, L., Isella, V., Guerini, F. R., Appollonio, I., Galli-Kienle, M., & Ferrarese, C. (2004). Increased susceptibility to plasma lipid peroxidation in Alzheimer disease patients. Curr Alzheimer Res, 1(2), 103–109. 10.2174/1567205043332171

Gevedon, O., Bolus, H., Lye, S. H., Schmitz, K., Fuentes-Gonzalez, J., Hatchell, K., Bley, L., Pienaar, J., Loewen, C., & Chtarbanova, S. (2019). In Vivo Forward Genetic Screen to Identify Novel Neuroprotective Genes in Drosophila melanogaster. J Vis Exp(149). 10.3791/59720

Gilestro, G. F., & Cirelli, C. (2009). pySolo: a complete suite for sleep analysis in Drosophila. Bioinformatics, 25(11), 1466–1467. 10.1093/bioinformatics/btp237

Gubb, D., Green, C., Huen, D., Coulson, D., Johnson, G., Tree, D., Collier, S., & Roote, J. (1999). The balance between isoforms of the Prickle LIM domain protein is critical for planar polarity in Drosophila imaginal discs. Genes & Development, 13(17), 2315–2327. DOI 10.1101/gad.13.17.2315

Guo, L., Zhang, N., & Simpson, J. H. (2022). Descending neurons coordinate anterior grooming behavior in Drosophila. Curr Biol, 32(4), 823–833 e824. 10.1016/j.cub.2021.12.055

Hahn, N., Geurten, B., Gurvich, A., Piepenbrock, D., Kastner, A., Zanini, D., Xing, G., Xie, W., Gopfert, M. C., Ehrenreich, H., & Heinrich, R. (2013). Monogenic heritable autism gene neuroligin impacts Drosophila social behaviour. Behav Brain Res, 252, 450–457. 10.1016/j.bbr.2013.06.020

Hoffman, T., Bar-Shalita, T., Granovsky, Y., Gal, E., Kalingel-Levi, M., Dori, Y., Buxbaum, C., Yarovinsky, N., & Weissman-Fogel, I. (2023). Indifference or hypersensitivity? Solving the riddle of the pain profile in individuals with autism. Pain, 164(4), 791–803. 10.1097/j.pain.0000000000002767

Homem, C. C., & Knoblich, J. A. (2012). Drosophila neuroblasts: a model for stem cell biology. Development, 139(23), 4297–4310. 10.1242/dev.080515

Hope, K. A., Flatten, D., Cavitch, P., May, B., Sutcliffe, J. S., O’Donnell, J., & Reiter, L. T. (2019). The Drosophila Gene Sulfateless Modulates Autism-Like Behaviors. Front Genet, 10, 574. 10.3389/fgene.2019.00574

Hotta, Y., & Benzer, S. (1970). Genetic dissection of the Drosophila nervous system by means of mosaics. Proc Natl Acad Sci U S A, 67(3), 1156–1163. 10.1073/pnas.67.3.1156

Hrycay, E. G., & Bandiera, S. M. (2015). Involvement of Cytochrome P450 in Reactive Oxygen Species Formation and Cancer. Adv Pharmacol, 74, 35–84. 10.1016/bs.apha.2015.03.003

Huai, J., & Zhang, Z. (2019). Structural Properties and Interaction Partners of Familial ALS-Associated SOD1 Mutants. Front Neurol, 10, 527. 10.3389/fneur.2019.00527

Huang, D. W., Sherman, B. T., & Lempicki, R. A. (2009a). Bioinformatics enrichment tools: paths toward the comprehensive functional analysis of large gene lists. Nucleic Acids Research, 37(1), 1–13. 10.1093/nar/gkn923

Huang, D. W., Sherman, B. T., & Lempicki, R. A. (2009b). Systematic and integrative analysis of large gene lists using DAVID bioinformatics resources. Nature Protocols, 4(1), 44–57. 10.1038/nprot.2008.211

Hussain, A., Pooryasin, A., Zhang, M., Loschek, L. F., La Fortezza, M., Friedrich, A. B., Blais, C. M., Ucpunar, H. K., Yepez, V. A., Lehmann, M., Gompel, N., Gagneur, J., Sigrist, S. J., & Grunwald Kadow, I. C. (2018). Inhibition of oxidative stress in cholinergic projection neurons fully rescues aging-associated olfactory circuit degeneration in Drosophila. Elife, 7. 10.7554/eLife.32018

Jin, B., Aung, T., Geng, Y., & Wang, S. (2020). Epilepsy and Its Interaction With Sleep and Circadian Rhythm. Front Neurol, 11, 327. 10.3389/fneur.2020.00327

Kanner, A. M. (2016). Management of psychiatric and neurological comorbidities in epilepsy. Nat Rev Neurol, 12(2), 106–116. 10.1038/nrneurol.2015.243

Katzenberger, R. J., Chtarbanova, S., Rimkus, S. A., Fischer, J. A., Kaur, G., Seppala, J. M., Swanson, L. C., Zajac, J. E., Ganetzky, B., & Wassarman, D. A. (2015). Death following traumatic brain injury in Drosophila is associated with intestinal barrier dysfunction. Elife, 4. 10.7554/eLife.04790

Katzenberger, R. J., Loewen, C. A., Wassarman, D. R., Petersen, A. J., Ganetzky, B., & Wassarman, D. A. (2013). A Drosophila model of closed head traumatic brain injury. Proc Natl Acad Sci U S A, 110(44), E4152–4159. 10.1073/pnas.1316895110

Kaye, A. D., Allen, K. E., Smith Iii, V. S., Tong, V. T., Mire, V. E., Nguyen, H., Lee, Z., Kouri, M., Jean Baptiste, C., Mosieri, C. N., Kaye, A. M., Varrassi, G., & Shekoohi, S. (2024). Emerging Treatments and Therapies for Autism Spectrum Disorder: A Narrative Review. Cureus, 16(7), e63671. 10.7759/cureus.63671

Khuong, T. M., Wang, Q. P., Manion, J., Oyston, L. J., Lau, M. T., Towler, H., Lin, Y. Q., & Neely, G. G. (2019). Nerve injury drives a heightened state of vigilance and neuropathic sensitization in Drosophila. Sci Adv, 5(7), eaaw4099. 10.1126/sciadv.aaw4099

King, L. B., Boto, T., Botero, V., Aviles, A. M., Jomsky, B. M., Joseph, C., Walker, J. A., & Tomchik, S. M. (2020). Developmental loss of neurofibromin across distributed neuronal circuits drives excessive grooming in Drosophila. PLoS Genet, 16(7), e1008920. 10.1371/journal.pgen.1008920

Kounatidis, I., Chtarbanova, S., Cao, Y., Hayne, M., Jayanth, D., Ganetzky, B., & Ligoxygakis, P. (2017). NF-kappaB Immunity in the Brain Determines Fly Lifespan in Healthy Aging and Age-Related Neurodegeneration. Cell Rep, 19(4), 836–848. 10.1016/j.celrep.2017.04.007

Kroll, J. R., Saras, A., & Tanouye, M. A. (2015). Drosophila sodium channel mutations: Contributions to seizure-susceptibility. Exp Neurol, 274(Pt A), 80–87. 10.1016/j.expneurol.2015.06.018

Lear, B. C., Lin, J. M., Keath, J. R., McGill, J. J., Raman, I. M., & Allada, R. (2005). The ion channel narrow abdomen is critical for neural output of the Drosophila circadian pacemaker. Neuron, 48(6), 965–976. 10.1016/j.neuron.2005.10.030

Li, J., Zhang, W., Guo, Z., Wu, S., Jan, L. Y., & Jan, Y. N. (2016). A Defensive Kicking Behavior in Response to Mechanical Stimuli Mediated by Drosophila Wing Margin Bristles. J Neurosci, 36(44), 11275–11282. 10.1523/JNEUROSCI.1416-16.2016

Li, W., Cressy, M., Qin, H., Fulga, T., Van Vactor, D., & Dubnau, J. (2013). MicroRNA-276a functions in ellipsoid body and mushroom body neurons for naive and conditioned olfactory avoidance in Drosophila. J Neurosci, 33(13), 5821–5833. 10.1523/JNEUROSCI.4004-12.2013

Lilienthal, A. J., Parida, M., & Manak, J. R. (2022). Characterization of prickle isoform-specific pk (pk1) and pk (sple1) mutations in Drosophila melanogaster. MicroPubl Biol, 2022. 10.17912/micropub.biology.000656

Lin, Y. J., Seroude, L., & Benzer, S. (1998). Extended life-span and stress resistance in the Drosophila mutant methuselah. Science, 282(5390), 943–946. 10.1126/science.282.5390.943

Lush, M. J., Li, Y., Read, D. J., Willis, A. C., & Glynn, P. (1998). Neuropathy target esterase and a homologous Drosophila neurodegeneration-associated mutant protein contain a novel domain conserved from bacteria to man. Biochem J, 332 ( *Pt 1*), 1–4. 10.1042/bj3320001

Madigan, J. P., Chotkowski, H. L., & Glaser, R. L. (2002). DNA double-strand break-induced phosphorylation of Drosophila histone variant H2Av helps prevent radiation-induced apoptosis. Nucleic Acids Res, 30(17), 3698–3705. 10.1093/nar/gkf496

Maitra, U., Scaglione, M. N., Chtarbanova, S., & O’Donnell, J. M. (2019). Innate immune responses to paraquat exposure in a Drosophila model of Parkinson’s disease. Sci Rep, 9(1), 12714. 10.1038/s41598-019-48977-6

Mapp, O. M., Walsh, G. S., Moens, C. B., Tada, M., & Prince, V. E. (2011). Zebrafish Prickle1b mediates facial branchiomotor neuron migration via a farnesylation-dependent nuclear activity. Development, 138(10), 2121–2132. 10.1242/dev.060442

Marcogliese, P. C., Deal, S. L., Andrews, J., Harnish, J. M., Bhavana, V. H., Graves, H. K., Jangam, S., Luo, X., Liu, N., Bei, D., Chao, Y. H., Hull, B., Lee, P. T., Pan, H., Bhadane, P., Huang, M. C., Longley, C. M., Chao, H. T., Chung, H. L.,…Yamamoto, S. (2022). Drosophila functional screening of de novo variants in autism uncovers damaging variants and facilitates discovery of rare neurodevelopmental diseases. Cell Rep, 38(11), 110517. 10.1016/j.celrep.2022.110517

Marsh, J. L., & Thompson, L. M. (2006). Drosophila in the study of neurodegenerative disease. Neuron, 52(1), 169–178. 10.1016/j.neuron.2006.09.025

Massingham, J. N., Baron, O., & Neely, G. G. (2021). Evaluating Baseline and Sensitised Heat Nociception in Adult Drosophila. Bio Protoc, 11(13), e4079. 10.21769/BioProtoc.4079

Matson, J. L., & Shoemaker, M. (2009). Intellectual disability and its relationship to autism spectrum disorders. Res Dev Disabil, 30(6), 1107–1114. 10.1016/j.ridd.2009.06.003

McDonald, N. M., Senturk, D., Scheffler, A., Brian, J. A., Carver, L. J., Charman, T., Chawarska, K., Curtin, S., Hertz-Piccioto, I., Jones, E. J. H., Klin, A., Landa, R., Messinger, D. S., Ozonoff, S., Stone, W. L., Tager-Flusberg, H., Webb, S. J., Young, G., Zwaigenbaum, L., & Jeste, S. S. (2020). Developmental Trajectories of Infants With Multiplex Family Risk for Autism: A Baby Siblings Research Consortium Study. JAMA Neurol, 77(1), 73–81. 10.1001/jamaneurol.2019.3341

McKenzie, K., Metcalfe, D., & Murray, A. L. (2023). Screening for intellectual disability in autistic people: A brief report. Research in Autism Spectrum Disorders, 100. ARTN 102076 10.1016/j.rasd.2022.102076

Mei, X., Wu, S., Bassuk, A. G., & Slusarski, D. C. (2013). Mechanisms of prickle1a function in zebrafish epilepsy and retinal neurogenesis. Dis Model Mech, 6(3), 679–688. 10.1242/dmm.010793

Min, K. T., & Benzer, S. (1999). Preventing neurodegeneration in the Drosophila mutant bubblegum. Science, 284(5422), 1985–1988. 10.1126/science.284.5422.1985

Moose, D. L., Haase, S. J., Aldrich, B. T., & Lear, B. C. (2017). The Narrow Abdomen Ion Channel Complex Is Highly Stable and Persists from Development into Adult Stages to Promote Behavioral Rhythmicity. Frontiers in Cellular Neuroscience, 11. ARTN 159 10.3389/fncel.2017.00159

Moraru, M. M., Egger, B., Bao, D. B., & Sprecher, S. G. (2012). Analysis of cell identity, morphology, apoptosis and mitotic activity in a primary neural cell culture system in Drosophila. Neural Dev, 7, 14. 10.1186/1749-8104-7-14

Moscato, E. H., Dubowy, C., Walker, J. A., & Kayser, M. S. (2020). Social Behavioral Deficits with Loss of Neurofibromin Emerge from Peripheral Chemosensory Neuron Dysfunction. Cell Rep, 32(1), 107856. 10.1016/j.celrep.2020.107856

Mrkusich, E. M., Flanagan, D. J., & Whitington, P. M. (2011). The core planar cell polarity gene prickle interacts with flamingo to promote sensory axon advance in the Drosophila embryo. Dev Biol, 358(1), 224–230. 10.1016/j.ydbio.2011.07.032

Mueller, J. M., Ravbar, P., Simpson, J. H., & Carlson, J. M. (2019). Drosophila melanogaster grooming possesses syntax with distinct rules at different temporal scales. PLoS Comput Biol, 15(6), e1007105. 10.1371/journal.pcbi.1007105

Myllymaki, H., & Ramet, M. (2014). JAK/STAT pathway in Drosophila immunity. Scand J Immunol, 79(6), 377–385. 10.1111/sji.12170

Neely, G. G., Keene, A. C., Duchek, P., Chang, E. C., Wang, Q. P., Aksoy, Y. A., Rosenzweig, M., Costigan, M., Woolf, C. J., Garrity, P. A., & Penninger, J. M. (2011). TrpA1 regulates thermal nociception in Drosophila. PLoS One, 6(8), e24343. 10.1371/journal.pone.0024343

Ng, J. (2012). Wnt/PCP proteins regulate stereotyped axon branch extension in Drosophila. Development, 139(1), 165–177. 10.1242/dev.068668

Nisar, S., Hashem, S., Bhat, A. A., Syed, N., Yadav, S., Azeem, M. W., Uddin, S., Bagga, P., Reddy, R., & Haris, M. (2019). Association of genes with phenotype in autism spectrum disorder. Aging (Albany NY), 11(22), 10742–10770. 10.18632/aging.102473

Niveditha, S., Ramesh, S. R., & Shivanandappa, T. (2017). Paraquat-Induced Movement Disorder in Relation to Oxidative Stress-Mediated Neurodegeneration in the Brain of Drosophila melanogaster. Neurochem Res, 42(11), 3310–3320. 10.1007/s11064-017-2373-y

Nukala, K. M., Lilienthal, A. J., Lye, S. H., Bassuk, A. G., Chtarbanova, S., & Manak, J. R. (2023). Downregulation of oxidative stress-mediated glial innate immune response suppresses seizures in a fly epilepsy model. Cell Rep, 42(1), 112004. 10.1016/j.celrep.2023.112004

Paemka, L., Mahajan, V. B., Skeie, J. M., Sowers, L. P., Ehaideb, S. N., Gonzalez-Alegre, P., Sasaoka, T., Tao, H., Miyagi, A., Ueno, N., Takao, K., Miyakawa, T., Wu, S., Darbro, B. W., Ferguson, P. J., Pieper, A. A., Britt, J. K., Wemmie, J. A., Rudd, D. S.,…Bassuk, A. G. (2013). PRICKLE1 interaction with SYNAPSIN I reveals a role in autism spectrum disorders. PLoS One, 8(12), e80737. 10.1371/journal.pone.0080737

Parker, L., Howlett, I. C., Rusan, Z. M., & Tanouye, M. A. (2011). Seizure and epilepsy: studies of seizure disorders in Drosophila. Int Rev Neurobiol, 99, 1–21. 10.1016/B978-0-12-387003-2.00001-X

Parker, L., Padilla, M., Du, Y., Dong, K., & Tanouye, M. A. (2011). Drosophila as a Model for Epilepsy: bss Is a Gain-of-Function Mutation in the Para Sodium Channel Gene That Leads to Seizures. Genetics, 187(2), 523–534. 10.1534/genetics.110.123299

Pessina, F., Gioia, U., Brandi, O., Farina, S., Ceccon, M., Francia, S., & d’Adda di Fagagna, F. (2021). DNA Damage Triggers a New Phase in Neurodegeneration. Trends Genet, 37(4), 337–354. 10.1016/j.tig.2020.09.006

Petersen, A. J., Katzenberger, R. J., & Wassarman, D. A. (2013). The innate immune response transcription factor relish is necessary for neurodegeneration in a Drosophila model of ataxia-telangiectasia. Genetics, 194(1), 133–142. 10.1534/genetics.113.150854

Petersen, A. J., Rimkus, S. A., & Wassarman, D. A. (2012). ATM kinase inhibition in glial cells activates the innate immune response and causes neurodegeneration in Drosophila. Proc Natl Acad Sci U S A, 109(11), E656–664. 10.1073/pnas.1110470109

Pfeiffenberger, C., Lear, B. C., Keegan, K. P., & Allada, R. (2010). Processing circadian data collected from the Drosophila Activity Monitoring (DAM) System. Cold Spring Harb Protoc, 2010(11), pdb prot5519. 10.1101/pdb.prot5519

Pinato, L., Galina Spilla, C. S., Markus, R. P., & da Silveira Cruz-Machado, S. (2019). Dysregulation of Circadian Rhythms in Autism Spectrum Disorders. Curr Pharm Des, 25(41), 4379–4393. 10.2174/1381612825666191102170450

Pinsonneault, R. L., Mayer, N., Mayer, F., Tegegn, N., & Bainton, R. J. (2011). Novel models for studying the blood-brain and blood-eye barriers in Drosophila. Methods Mol Biol, 686, 357–369. 10.1007/978-1-60761-938-3_17

Puttachary, S., Sharma, S., Stark, S., & Thippeswamy, T. (2015). Seizure-induced oxidative stress in temporal lobe epilepsy. Biomed Res Int, 2015, 745613. 10.1155/2015/745613

Qian, H., Shao, M., Wei, Z., Zhang, Y., Liu, S., Chen, L., & Meng, J. (2024). Intact painful sensation but enhanced non-painful sensation in individuals with autistic traits. Front Psychiatry, 15, 1432149. 10.3389/fpsyt.2024.1432149

Qiu, S., Qiu, Y., Li, Y., & Cong, X. (2022). Genetics of autism spectrum disorder: an umbrella review of systematic reviews and meta-analyses. Transl Psychiatry, 12(1), 249. 10.1038/s41398-022-02009-6

Ravbar, P., Zhang, N., & Simpson, J. H. (2021). Behavioral evidence for nested central pattern generator control of Drosophila grooming. Elife, 10. 10.7554/eLife.71508

Samuels, T. J., Arava, Y., Jarvelin, A. I., Robertson, F., Lee, J. Y., Yang, L., Yang, C. P., Lee, T., Ish-Horowicz, D., & Davis, I. (2020). Neuronal upregulation of Prospero protein is driven by alternative mRNA polyadenylation and Syncrip-mediated mRNA stabilisation. Biol Open, 9(5). 10.1242/bio.049684

Santana, J. F., Parida, M., Long, A., Wankum, J., Lilienthal, A. J., Nukala, K. M., & Manak, J. R. (2020). The Dm-Myb Oncoprotein Contributes to Insulator Function and Stabilizes Repressive H3K27me3 PcG Domains. Cell Rep, 30(10), 3218–3228 e3215. 10.1016/j.celrep.2020.02.053

Schneider, C. A., Rasband, W. S., & Eliceiri, K. W. (2012). NIH Image to ImageJ: 25 years of image analysis. Nat Methods, 9(7), 671–675. 10.1038/nmeth.2089

Seluzicki, A., Flourakis, M., Kula-Eversole, E., Zhang, L., Kilman, V., & Allada, R. (2014). Dual PDF signaling pathways reset clocks via TIMELESS and acutely excite target neurons to control circadian behavior. PLoS Biol, 12(3), e1001810. 10.1371/journal.pbio.1001810

Shadfar, S., Parakh, S., Jamali, M. S., & Atkin, J. D. (2023). Redox dysregulation as a driver for DNA damage and its relationship to neurodegenerative diseases. Transl Neurodegener, 12(1), 18. 10.1186/s40035-023-00350-4

Sharp, K. A., & Axelrod, J. D. (2016). Prickle isoforms control the direction of tissue polarity by microtubule independent and dependent mechanisms. Biol Open, 5(3), 229–236. 10.1242/bio.016162

Shimizu, K., Sato, M., & Tabata, T. (2011). The Wnt5/planar cell polarity pathway regulates axonal development of the Drosophila mushroom body neuron. J Neurosci, 31(13), 4944–4954. 10.1523/JNEUROSCI.0154-11.2011

Song, Y., Zhang, X., Wang, B., Luo, X., Zhang, K., Zhang, X., Wu, Q., & Sun, M. (2024). BPAP induces autism-like behavior by affecting the expression of neurodevelopmental genes in Drosophila melanogaster. Ecotoxicol Environ Saf, 288, 117405. 10.1016/j.ecoenv.2024.117405

Sowers, L. P., Loo, L., Wu, Y., Campbell, E., Ulrich, J. D., Wu, S., Paemka, L., Wassink, T., Meyer, K., Bing, X., El-Shanti, H., Usachev, Y. M., Ueno, N., Manak, J. R., Shepherd, A. J., Ferguson, P. J., Darbro, B. W., Richerson, G. B., Mohapatra, D. P.,…Bassuk, A. G. (2013). Disruption of the non-canonical Wnt gene PRICKLE2 leads to autism-like behaviors with evidence for hippocampal synaptic dysfunction. Mol Psychiatry, 18(10), 1077–1089. 10.1038/mp.2013.71

Sridhar, V. H., Roche, D. G., & Gingins, S. (2019). Tracktor: Image-based automated tracking of animal movement and behaviour. Methods in Ecology and Evolution, 10(6), 815–820. 10.1111/2041-210x.13166

Stahl, A., & Tomchik, S. M. (2024). Modeling neurodegenerative and neurodevelopmental disorders in the Drosophila mushroom body. Learn Mem, 31(5). 10.1101/lm.053816.123

Stojkovic, M., Petrovic, M., Capovilla, M., Milojevic, S., Makevic, V., Budimirovic, D. B., Corscadden, L., He, S., & Protic, D. (2024). Using a Combination of Novel Research Tools to Understand Social Interaction in the Drosophila melanogaster Model for Fragile X Syndrome. Biology (Basel), 13(6). 10.3390/biology13060432

Stone, M. C., Roegiers, F., & Rolls, M. M. (2008). Microtubules have opposite orientation in axons and dendrites of Drosophila neurons. Mol Biol Cell, 19(10), 4122–4129. 10.1091/mbc.e07-10-1079

Szebenyi, A. L. (1969). Cleaning Behaviour in Drosophila-Melanogaster. Animal Behaviour, 17, 641–&. Doi 10.1016/S0003-3472(69)80006-0

Tao, H., Manak, J. R., Sowers, L., Mei, X., Kiyonari, H., Abe, T., Dahdaleh, N. S., Yang, T., Wu, S., Chen, S., Fox, M. H., Gurnett, C., Montine, T., Bird, T., Shaffer, L. G., Rosenfeld, J. A., McConnell, J., Madan-Khetarpal, S., Berry-Kravis, E.,…Bassuk, A. G. (2011). Mutations in prickle orthologs cause seizures in flies, mice, and humans. Am J Hum Genet, 88(2), 138–149. 10.1016/j.ajhg.2010.12.012

Tener, S. J., Lin, Z., Park, S. J., Oraedu, K., Ulgherait, M., Van Beek, E., Martinez-Muniz, A., Pantalia, M., Gatto, J. A., Volpi, J., Stavropoulos, N., Ja, W. W., Canman, J. C., & Shirasu-Hiza, M. (2024). Neuronal knockdown of Cullin3 as a Drosophila model of autism spectrum disorder. Sci Rep, 14(1), 1541. 10.1038/s41598-024-51657-9

Tomchik, S. M., & Davis, R. L. (2009). Dynamics of learning-related cAMP signaling and stimulus integration in the Drosophila olfactory pathway. Neuron, 64(4), 510–521. 10.1016/j.neuron.2009.09.029

Tully, T., & Quinn, W. G. (1985). Classical conditioning and retention in normal and mutant Drosophila melanogaster. J Comp Physiol A, 157(2), 263–277. 10.1007/BF01350033

Viscidi, E. W., Triche, E. W., Pescosolido, M. F., McLean, R. L., Joseph, R. M., Spence, S. J., & Morrow, E. M. (2013). Clinical characteristics of children with autism spectrum disorder and co-occurring epilepsy. PLoS One, 8(7), e67797. 10.1371/journal.pone.0067797

Wang, H., Lautrup, S., Caponio, D., Zhang, J., & Fang, E. F. (2021). DNA Damage-Induced Neurodegeneration in Accelerated Ageing and Alzheimer’s Disease. Int J Mol Sci, 22(13). 10.3390/ijms22136748

Wang, M., Zhang, X., Zhong, L., Zeng, L., Li, L., & Yao, P. (2025). Understanding autism: Causes, diagnosis, and advancing therapies. Brain Res Bull, 227, 111411. 10.1016/j.brainresbull.2025.111411

Wang, Y., Chiang, A. S., Xia, S., Kitamoto, T., Tully, T., & Zhong, Y. (2003). Blockade of neurotransmission in Drosophila mushroom bodies impairs odor attraction, but not repulsion. Curr Biol, 13(21), 1900–1904. 10.1016/j.cub.2003.10.003

Wise, A., Tenezaca, L., Fernandez, R. W., Schatoff, E., Flores, J., Ueda, A., Zhong, X., Wu, C. F., Simon, A. F., & Venkatesh, T. (2015). Drosophila mutants of the autism candidate gene neurobeachin (rugose) exhibit neuro-developmental disorders, aberrant synaptic properties, altered locomotion, and impaired adult social behavior and activity patterns. J Neurogenet, 29(2-3), 135–143. 10.3109/01677063.2015.1064916

Xavier, S. D. (2021). The relationship between autism spectrum disorder and sleep. Sleep Sci, 14(3), 193–195. 10.5935/1984-0063.20210050

Yang, T., Bassuk, A. G., Stricker, S., & Fritzsch, B. (2014). Prickle1 is necessary for the caudal migration of murine facial branchiomotor neurons. Cell Tissue Res, 357(3), 549–561. 10.1007/s00441-014-1925-6

Yellman, C., Tao, H., He, B., & Hirsh, J. (1997). Conserved and sexually dimorphic behavioral responses to biogenic amines in decapitated Drosophila. Proc Natl Acad Sci U S A, 94(8), 4131–4136. 10.1073/pnas.94.8.4131

Young, A. P., Jackson, D. J., & Wyeth, R. C. (2020). A technical review and guide to RNA fluorescence in situ hybridization. PeerJ, 8, e8806. 10.7717/peerj.8806

Younger, M. A., Herre, M., Goldman, O. V., Lu, T.-C., Caballero-Vidal, G., Qi, Y., Gilbert, Z. N., Gong, Z., Morita, T., Rahiel, S., Ghaninia, M., Ignell, R., Matthews, B. J., Li, H., & Vosshall, L. B. (2022). Non-Canonical Odor Coding in the Mosquito. bioRxiv, 2020.2011.2007.368720. 10.1101/2020.11.07.368720

Zahra, A., Wang, Y., Wang, Q., & Wu, J. (2022). Shared Etiology in Autism Spectrum Disorder and Epilepsy with Functional Disability. Behav Neurol, 2022, 5893519. 10.1155/2022/5893519

