## Supplementary Figures 1-7 for "*Drosophila Prickle* Mutants Display Comorbid Neurological Phenotypes and Provide A Genetic Link Between Epilepsy and Autism Spectrum Disorder"

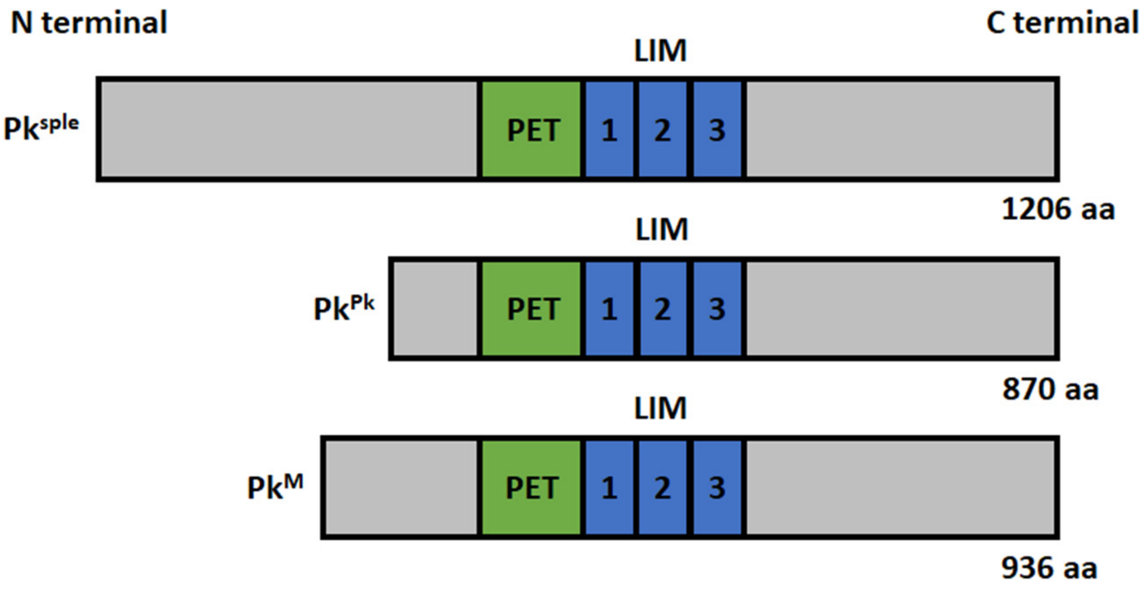
N terminal
C terminal
LIM
Pk^sple
PET
1
2
3
1206 aa
LIM
Pk^Pk
PET
1
2
3
870 aa
LIM
Pk^M
PET
1
2
3
936 aa

**Supplementary Figure 1. Schematic of the adult Prickle protein isoforms, $Pk^{sple}$, $Pk^{pk}$ and $Pk^{M}$.** The three adult protein isoforms of Prickle share one PET and three LIM domains while the amino-terminal ends differentiate the three isoforms.

pk-pk: AGUCUGCUGCGCGCCUCGCACGAACAUCGAACUCGCGGACGUGCGC
GUGCGCGAACGGAACGCUCGAAGGCAAACGAAACGGAAAGGAACGUUAGG
GAACGCACGGAAACGGAACGGAUCAGCGACGGCCCCCGCGUUUUUGCUUU
UUUGCGCUUUUGGGCUUAUCAAGUGCCGGGAAUCGGUGAACGGGAGUCCA
GAAAUCGGGCAGCUGUCGCGUGCAGUGGCCAGGAAAAGGAGUUUGCGGAA
AAUAAUGAAAUGCCUGCCGUCGGUGGCAGCAGUGUGGCUGCGUGUGUUUC
UUGAGCGGCCGGGAAAUGUGCAUCAAUUAUUUUAAAUGGCCAGCAGCCACC
CCCUCCUCACCCAACCCCUGCGAAGUGUCCCCCACCCCAAGCACAUGCAUA
AAUACAAUUAGGCAGCUAGAUCGGGCCUGAGCCAUUUGGUGGAGCCAAAGU
UAAUUCGGUGCGCGAACCAGUUUAUCGACAGGCACUCCGCGAAUUUUUUCA
CGAACGCGCCAAGAAAUUUAUAAUAAAUACGCCUAAGCUCAAAUUGUUGUCA
GCUGGCUCGGCUUUUGUUGAUUUAUGUGCGCGAUUAAAGGAAACAACGCAG
CGCAGUCGGCCGAGAAACCGCAGAAUAAGAUAAUAAAUAUAAGUAAUGCCG
AAAGGCAUCACCCCCGCCCCCAGUGUCACCAGUGAAAUACCUGUGAGUGCC
AAUCAAGCGACAAAGAAACCGAAAAGACAUCAGCCCGUAGAACGCCAGCCAU
AAAAACUUCGGAUACAUCCUCCGCUCCGCAAACAUGGAUACCCCAAAUCAAA
UGCCUGUUGAGCUAGAAAG

pk-sple:
CAAUGGUGGCGCGUCUGCUUCUUGUACGGGGACCAGCAGAAAUAUUAUCGA
CAGCUCUAUAGCAAGGCGGCGGCCCAGCGGCUGGCGGACGCUAAUCAGGA
ACCCGACAAUGCCCGCGAUCGGGAGUACGACACCGUGGACUGCGAUCUUAU
CGCCGGGCAAUUAGAUGCCGUCGAGGAUGCGGACGAUGGCAUCGAUCUAG
GCGAUCACUCGUCGACCCCGAAGGGAGGUGCGACGACGGCGGGCCGUCCG
CUCUUCCCGCAUUCCUCCUCACCGCGGCGCAGCAAGAAGCUCCUGAGAUCC
CUGCGAGCCCAUGUGCGCGGGGAGAAGCUGCCGAAGAACGACACUACGAC
GGCUAACGAGUCCAGCGAGGUGACCCAGCGGAAUGCCAGGGUGACCGUUC
UCGACGACCCCUUUCUAUUCGGCAUCGACGCCGAUCACUUGGGCGAUCUC
GUGGUGCGGGGCAAGCGGUACAGUACGCUAGACGCCACCGAGAACAUGGC
CAGGUUCUAUGCGGAGCAGGAGGCCACGGCCCAGGUCCUGGAGAUCAUCG
AGCAGGAAGAAGAAUCUCCCGAGCAGGAAGCUCCCAAGCCCGCCCUACCGC
CCAAACAGAAGCAGCAGCGUCCUGUGCCGCCACUGCCACCACCACCAGCGA
ACCGGGUCACCCAGGACCAAGGAACGCAACCUGCAGCACCACAAGUGCCCC
UGCAACCCCUGACCGCCGGCGAUCUUCAGUUUCUAAACUUGAGCCUGCGGC
AGAGGAGUUUACCGCGCAGCAUGAAGCCGUUUAAGGAUGCCCACGAUAUCA
GUUUCACUUUCAACGAACUGGACACGAGCGCCGAACCGGAAGUGGCGACAG
GAGCCGCCCAGCAGGAGUCAAAUGA

pk-M:
AAGAACCAUCCGUGAGUGCAACGCGAGCGGACGGAUCCGCCGAUUCCUCGU
GUAUUCCGACUUAUCCCGCUGCUGAUAAGUCGCCGCGAUUGCAACAAUUAG
CUAUAAUUAUUCAUUAAUUCGAAUCGGUGUCGAGGAGUUAUCGCUAAUUCG
CAGAAACUGUGCCAGCGAGAUAGAAUACCACUCAAGUUUUCCUCAAUCUCU
UCGCCGGUCGCCUUUGUUAUUGUUUGCCGCUGCUUCUUCCUCUUUUUUG
GGGGGAAUAGUUCUUUGUGUUUGUGCCCGUGUGUUUGAGAGCCUGGCAGU
GUGUGUGUACGUAUCUGUGUCUGUGAGUCAGCGCAAAUGGCAAUGCGCAU
CUUUGUUUACCCGCAAAAUCCAGGGCUCUCAUCGCGGAGAAGUGCCAGUGA
AAGCGGCGCCAAAAUCUUAGUCUAGUGAUUUUCUUCUGCCUUUCCGGCCGA
AACAAUGAACGAUUCGACCGAUAAUUUGCAUGCCGACUGCGACGGUCGAGU
CAGCAACAACAACAAUGGCAACAGCAACACCAACGAUGGACCCAACAACGAC
GGCGACUCCGACGAGGAGGUCAUCGAGGGCAUGGCGCUGCUGGAGGGCAA
CUACCAAGUGCUGCGCCAGUGGGUGCCGCCCGCCCCCAACUAUUGGGAUG
CGCCCCCCAAGGCCAUCAUCAAAUCCGCCGAAGUCAG

**Supplementary Figure 2.** ***prickle*** **isoform probe sequences for** ***in situ*** **hybridization.**

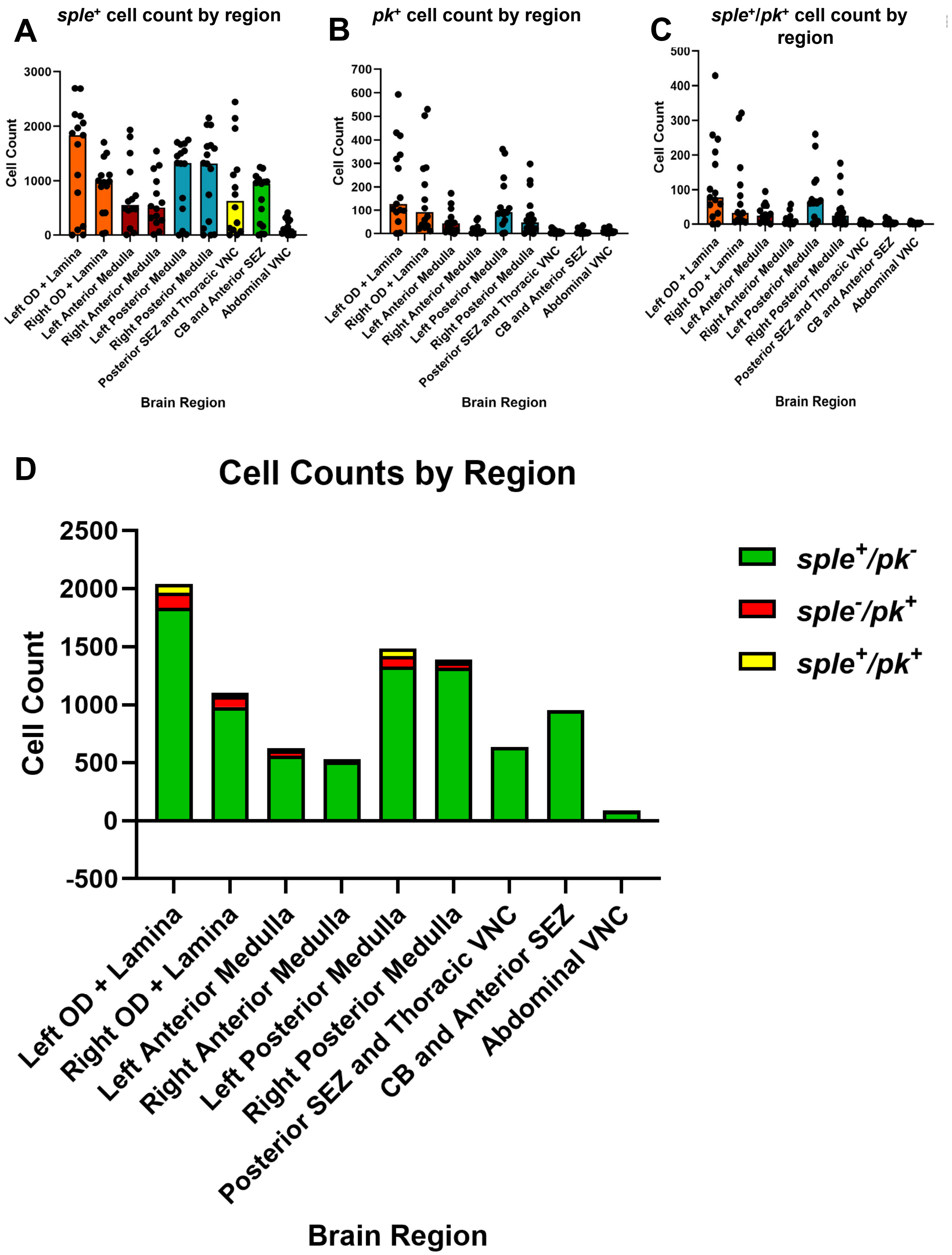
A
sple+ cell count by region
Cell Count
Brain Region
B
pk+ cell count by region
C
sple+/pk+ cell count by region
Left OD + Lamina
Right OD + Lamina
Left Anterior Medulla
Right Anterior Medulla
Left Posterior Medulla
Right Posterior Medulla
Posterior SEZ and Thoracic VNC
CB and Anterior SEZ
Abdominal VNC
D
Cell Counts by Region
sple+/pk-
sple-/pk+
sple+/pk+

**Supplementary Figure 3. Wild-type third-instar larval brains show widespread *$pk^{sple}$* isoform expression and limited *$pk^{pk}$* isoform expression.** Each data point represents the number of cells expressing the A) *$pk^{sple}$* and B) *$pk^{pk}$* transcript isoforms in each slice through the brain. Data shown is mean cell count ± SEM, n ≥ 12 1-um slices for each region. C) The number of *$pk^{sple}$*+ cells also expressing *$pk^{pk}$* in each region. Data shown is mean cell count ± SEM, n ≥ 12 1-um slices for each region. D) Number of cells expressing each combination of isoforms in each region of the brain. Graph bars represent the mean number of cells expressing either *$pk^{sple}$+/$pk^{pk}$-* (green), *$pk^{sple}$-/$pk^{pk}$+*(red), or *$pk^{sple}$+/$pk^{pk}$+*(yellow) in each region of the brain.

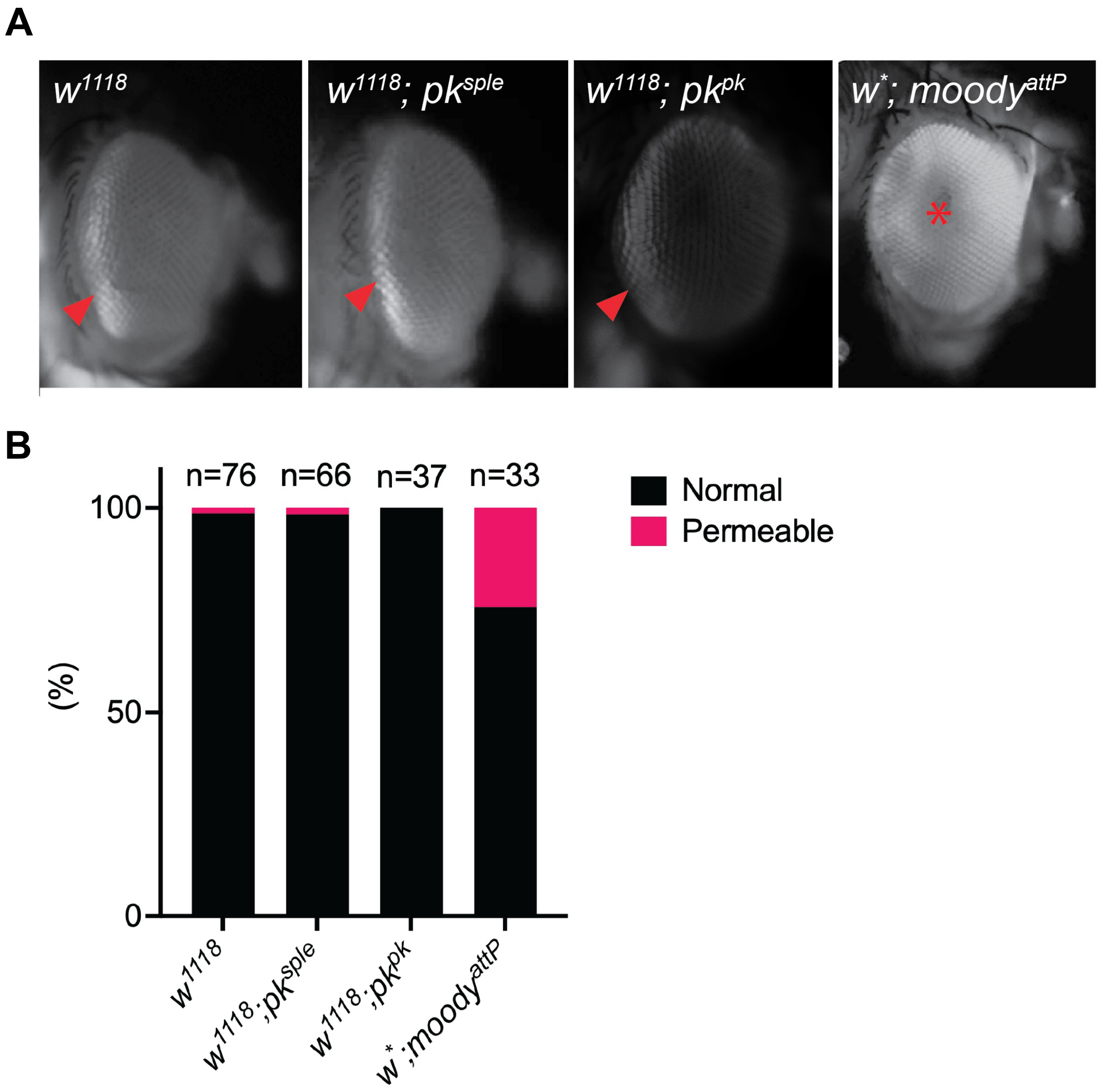
A
w1118
w1118; pksple
w1118; pkpk
w*; moodyattP
B
n=76
n=66
n=37
n=33
100
50
0
(%)
w1118
w1118;pksple
w1118;pkpk
w*;moodyattP
Normal
Permeable

**Supplementary Figure 4. The blood brain barrier is intact in *pk^pk* flies. (A)** Representative images of eyes of +/+ (*$w^{1118}$*), *sple/sple* (*$w^{1118}$; $pk^{sple}$*), *pk/pk* ($w^{1118}$; $pk^{pk}$) and *$moody^{attP}$/ $moody^{attP}$* (*$w^{*}$; $moody^{attP,}$* positive control) female flies (age range 3-10 days-old) injected with 10,000 MW Alexa488-Dextran. The accumulation of fluorescence at the border of the eye (red arrowhead) corresponds to the hemolymph exclusion line which reflects an intact Blood Eye Barrier (BEB). BEB is used as a proxy for Blood Brain Barrier (BBB) integrity. Diffused eye fluorescence indicating permeable BEB/BBB (red asterisk) is observed in moody mutants. **(B)** Quantification of BEB integrity in indicated genotypes based on the presence of hemolymph exclusion line (Normal BBB) or diffused fluorescence (Permeable BBB). Data is graphed as a percentage of flies having normal or permeable BBB; n indicates sample size for each genotype.

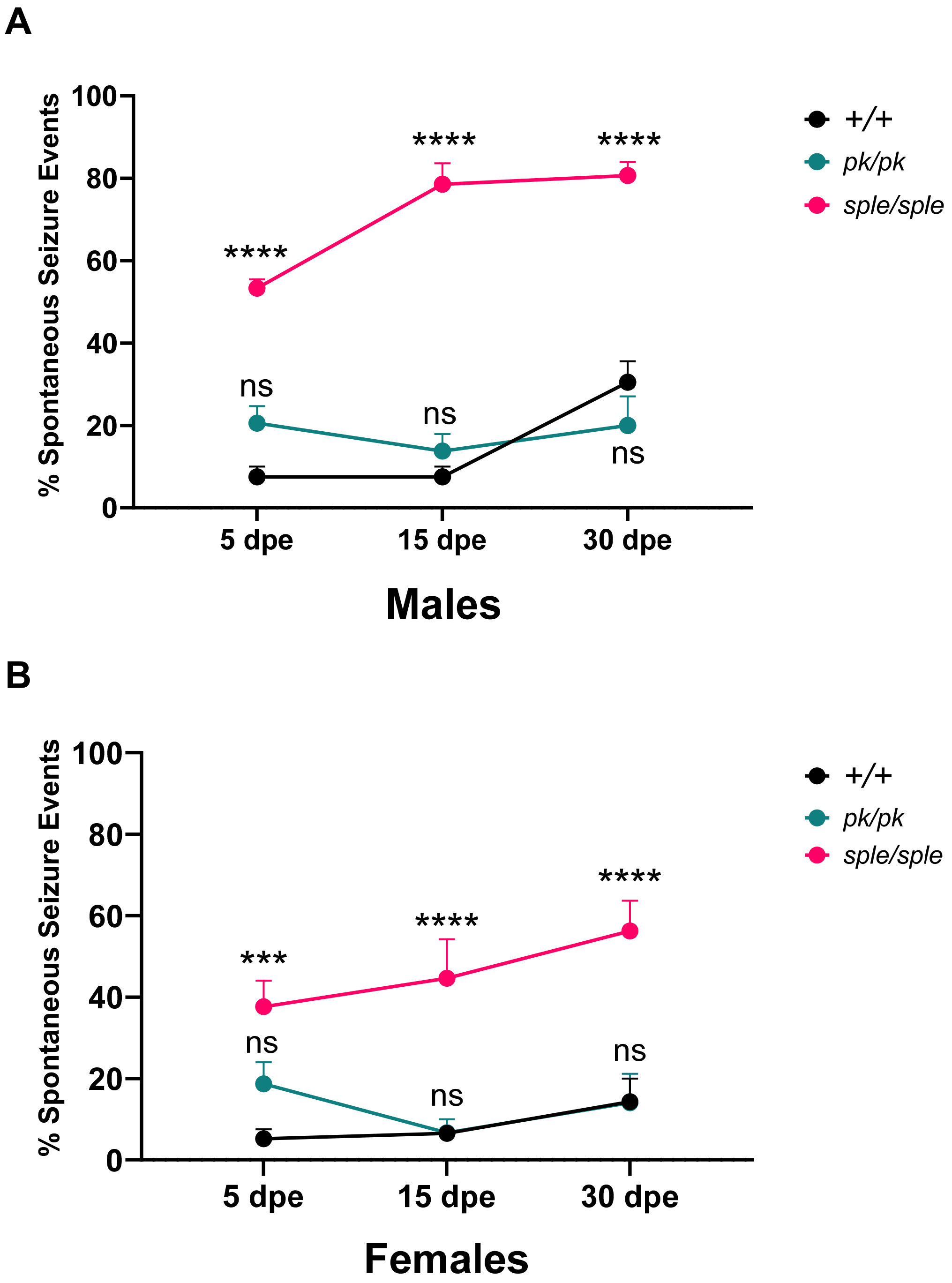
A
% Spontaneous Seizure Events
100
80
60
40
20
0
5 dpe
15 dpe
30 dpe
Males
****
****
****
ns
ns
ns
+/+
pk/pk
sple/sple
B
% Spontaneous Seizure Events
100
80
60
40
20
0
5 dpe
15 dpe
30 dpe
Females
***
****
****
ns
ns
ns
+/+
pk/pk
sple/sple

**Supplementary Figure 5. Homozygous *pk*$^{sple}$ but not *pk*$^{pk}$ mutants display increased percentage of spontaneous seizure events. A,** Quantification of percentage of spontaneous seizure events of 5, 15 and 30 day old male +/+, *sple/sple* and *pk/pk* mutants. Male *sple/sple* mutants show significant increase at all timepoints while there is no significant difference in the percentage of spontaneous seizure events of *pk/pk* mutants when compared to controls across all timepoints. Data are Mean ± SEM analyzed by a Two-way ANOVA with Tukey's multiple comparisons test. n = 4-8 replicates (8-10 files per replicate) for all genotypes and timepoints. **B,** Quantification of percentage of spontaneous seizure events of 5,15, and 30 day old female +/+, *sple/sple* and *pk/pk* mutants. Female *sple/sple* mutants show significant increase at all timepoints while there is no significant difference in the percentage of spontaneous seizure events of *pk/pk* mutants when compared to controls across all timepoints. Data are Mean ± SEM analyzed by a Two-way ANOVA with Tukey's multiple comparisons test. n = 3-8 replicates (8-10 files per replicate) for all genotypes and timepoints.

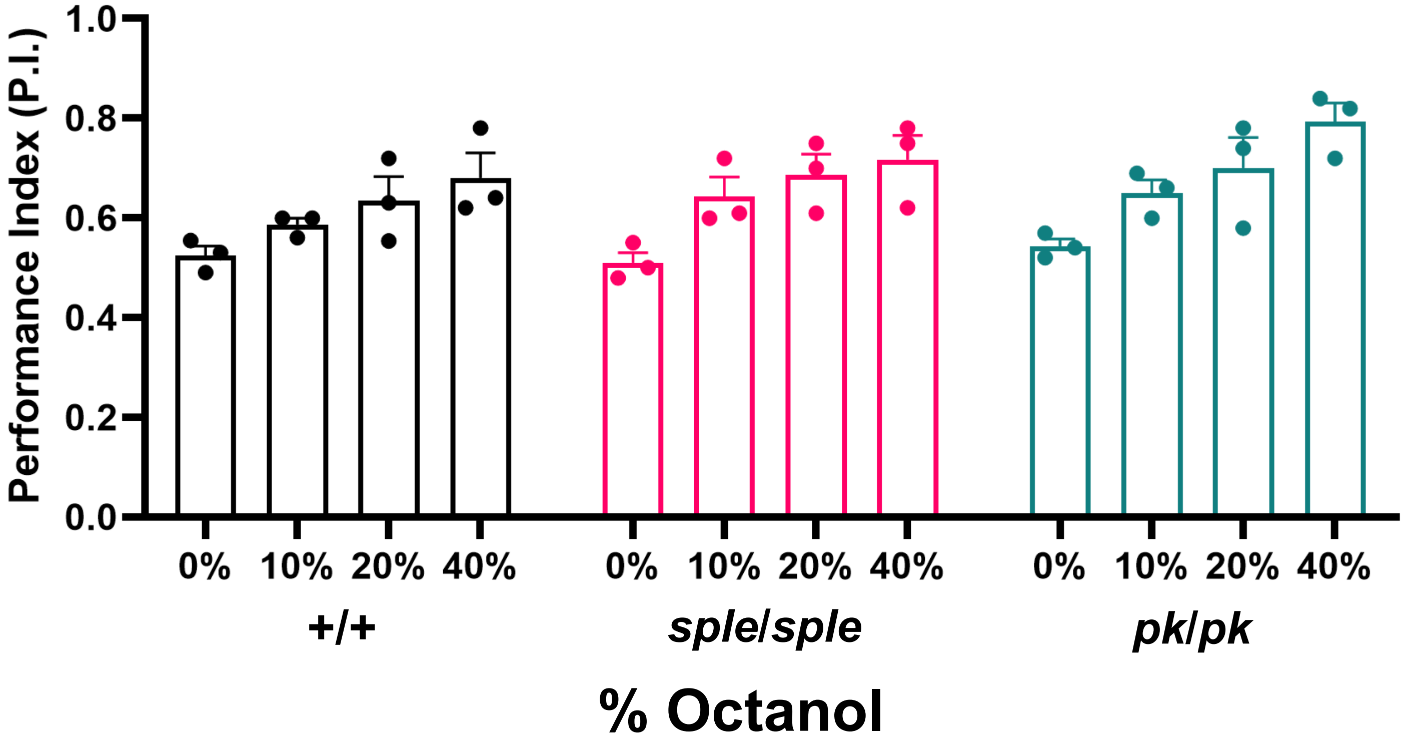

Performance Index (P.I.)
1.0
0.8
0.6
0.4
0.2
0.0
0%
10%
20%
40%
+/+
*sple/sple*
*pk/pk*
% Octanol

**Supplementary Figure 6. Both Homozygous *pk*$^{sple}$ and *pk*$^{pk}$ mutants flies can perceive odors.** Olfactory aversion assay demonstrates that *pk/pk* and *sple/sple* mutants are able to perceive and avoid increasing concentrations of 3-octanol comparable to controls. n = 3 biological replicates with 50 flies per replicate for all genotypes.

| Phenotype | | Presence of Defect | |
|---|---|---|---|
| | | *sple/sple* | *pk/pk* |
| **Innate Immune Response Upregulation** | | ✓ | ✓ |
| **Redox Pathway Gene Upregulation** | | ✓ | ✓ |
| **Blood Brain Barrier Breach** | | - | - |
| **Neuron Loss** | Sustained Neuronal Cell Death | + | ++ |
| | High Neurodegeneration Index | - | ✓ |
| **DNA Breaks** | | - | ✓ |
| **Reduced Lifespan** | | + | ++ |
| **Locomotor Defects** | Spontaneous Seizures | ✓ | - |
| | Increased Median Turning Angle | ✓ | - |
| | Reduced Distance Travelled | ++ | + |
| | Pronounced Climbing Defects | ✓ | - |
| **Increased Nociceptive Response** | | ✓ | - |
| **Learning/Memory Defects** | Olfaction Response Defect | - | - |
| | Learning Disability | ✓ | - |
| **Social Isolation** | Median Interfly Distance | ✓ | - |
| **Communication** | Decreased courtship behavior | ✓ | - |
| **Repetitive Behavior** | Increased grooming | +++ | - |
| **Circadian Rhythm & Visualization Defects** | Average AM Anticipation Defect | - | ✓ |
| | Average PM Anticipation Defect | - | ✓ |
| | Reduced Rhythmicity | ✓ | ✓ |
| | Decreased Depolarization Amplitude (eye) | - | - |

**Supplementary Figure 7. Summary of phenotypes in Homozygous *pk*$^{sple}$ and *pk*$^{pk}$ mutants.**
Check mark indicates “yes”, “-” indicates “no”; level of severity of phenotypes is indicated by “+”.
